# Multi-omics profiling with indoor-unmanned phenotyping reveals drought adaptation through constitutive *ABF1* expression in wild rice

**DOI:** 10.1101/2025.09.08.674796

**Authors:** Fumiyuki Soma, Yoshihiro Kawahara, Takanari Tanabata, Atsushi Hayashi, Yuka Kitomi, Nobuo Kochi, Eiji Yamamoto, Nobuhiro Tanaka, Michiya Negishi, Kenichi Tokuda, Hiroaki Sakai, Yusaku Uga

## Abstract

Improving drought resistance is crucial for stable crop production under climate change. Identifying the mechanisms of drought resistance using diverse genetic resources, including crop wild relatives, would be beneficial for molecular breeding. Here, we developed an indoor, unmanned phenotyping platform that can noninvasively and automatically collect temporal data on plant responses to drought stress. Using this system, we analyzed the phenotypic, transcriptomic, and environmental data of four cultivated rice varieties and five wild relatives. Multi-omics analysis revealed that one wild rice species exhibited drought adaptation through the constitutive expression of *ABSCISIC ACID RESPONSIVE ELEMENT-BINDING FACTOR 1* (*ABF1*), which encodes a transcription factor that regulates drought resistance, before drought stress. Drought testing of introgression lines of cultivated rice with constitutive *ABF1* expression revealed higher drought tolerance than in cultivars without a growth penalty. Our findings suggest that constitutive *ABF1* expression contributes to drought adaptation in both cultivated and wild rice.

---

Drought stress is a major factor limiting crop yields^1,2^. Developing high-yield drought-resistant crops is vital for ensuring global food security. Biological analyses using model plants, such as *Arabidopsis,* have revealed the basic molecular mechanisms of the drought response^2,3^. The drought resistance strategies of plants are classified into three major mechanisms: avoidance, escape, and tolerance^4^. These mechanisms are primarily regulated by the plant hormone abscisic acid (ABA). Under drought conditions, ABA accumulation within the plant promotes stomatal closure, resulting in reduced water loss^5–7^. Additionally, ABA induces drought-responsive genes encoding functional proteins, such as dehydrins and late embryogenesis-abundant proteins, to protect plant cells from damage in response to drought^8^. The bZIP transcription factor ABA RESPONSIVE ELEMENT-BINDING FACTOR 1 (ABF1/bZIP12) acts as a central regulator of ABA signaling in rice^9,10^. ABA accumulation activates *ABF1* expression, inducing drought stress-responsive genes in rice. *ABF1* overexpressing rice lines exhibit higher drought tolerance^10^. The DEHYDRATION-RESPONSIVE ELEMENT-BINDING 2A (DREB2A) transcription factor, whose expression is induced later than that of *ABF1*, acquires substantial drought tolerance in an ABA-independent manner in rice^11,12^. Applying these findings to other crops could lead to the development of drought-resistant crops. However, overexpression of stress-responsive genes often causes growth retardation, leading to yield penalties^13^. Therefore, achieving finely tuned regulation of drought-responsive gene expression is essential for developing drought-tolerant crops while maintaining normal growth and productivity.

Cultivated rice (*Oryza sativa* L.) is a staple food worldwide but is sensitive to drought stress^3^. One reason could be the tradeoff between improved grain yield and stress tolerance loss that occurred during domestication^14^. Conversely, wild relatives may serve as valuable genetic resources for stress tolerance traits^15–18^. Introducing drought-resistant genes from wild relatives may enable cultivated rice to withstand yield reductions due to drought stress^17^. For example, the elite *DROUGHT1* allele, which enhances drought tolerance by adjusting cell wall structure, is considered to have originated from *O. rufipogon*, the wild progenitor of Asian cultivated rice^19^. Similarly, the *O. rufipogon-* derived *LEA12* allele improves drought and salt tolerance^20^. Promising alleles from wild rice offer potential for improving stress tolerance in cultivated rice.

Several wild rice species exhibit drought resistance superior to that of cultivated rice. Hybrid offspring derived from crosses between wild and cultivated rice have demonstrated drought tolerance^15,21^. However, identifying drought-resistance genes from natural variations via QTL mapping or genome-wide association studies is time-consuming, labor-intensive, and requires large sample populations. In contrast, multi-omics approaches, integrating data from transcriptomics, proteomics, and other omics, offer a comprehensive strategy for identifying genes involved in plant physiological responses^22,23^. Recent multi-omics approaches using wild and cultivated rice have elucidated the underlying mechanisms of metabolic pathways and drought stress responses^12,24^. Thus, a comparative multi-omics analysis of cultivated and wild rice is expected to identify drought-adaptation mechanisms unique to wild rice.

Many agronomic traits are typically assessed in the field. However, when investigating drought resistance, researchers simulate drought conditions using tools such as rainout shelters and irrigation systems. Despite these efforts, effectively assessing stress responses in the field remains a significant challenge owing to various factors, including weather, inconsistent management practices, and other abiotic and biotic stresses. These complicate the evaluation of targeted stress responses in field trials. Plant phenotyping platforms that replicate field conditions have been developed^25–27^. For example, PhenoSphere, a large-scale cultivation platform with controlled environmental conditions, simulates field-like environments^26^. The agro-environment emulator enables plant growth evaluation under artificial environmental conditions that mimic predicted future global warming^27^. Previously, we developed an Internet of Things (IoT)-based pot system for controlled soil water treatments (iPOTs) that can simulate drought conditions by controlling soil water levels^12,25,28^. However, to detect subtle morphological and physiological responses to drought stress, existing platforms should incorporate new noninvasive and automated plant phenotyping capabilities.

This study aimed to improve drought resistance in rice by identifying the mechanisms of drought resistance using diverse genetic resources, including wild relatives. We constructed an unmanned phenotyping platform in growth chambers that combined a movable multi-camera system with a modified automatic irrigation system, iPOTs, to automatically obtain temporal data on plant behavior in response to drought stress.

## Results

### Unmanned phenotyping platform development

We developed an unmanned phenotyping platform designated IoT-based Platform of Unmanned Phenotyping with Imitated Land condition (iPUPIL). Each growth chamber was equipped with three key systems: movable multi-camera, automatic irrigation, and environmental sensor systems (Fig. 1, Supplementary Fig. 1). These iPUPIL functions enable automatic recording of images that capture morphological and physiological states, as well as the environmental conditions of individual plants.

**Figure 1.**
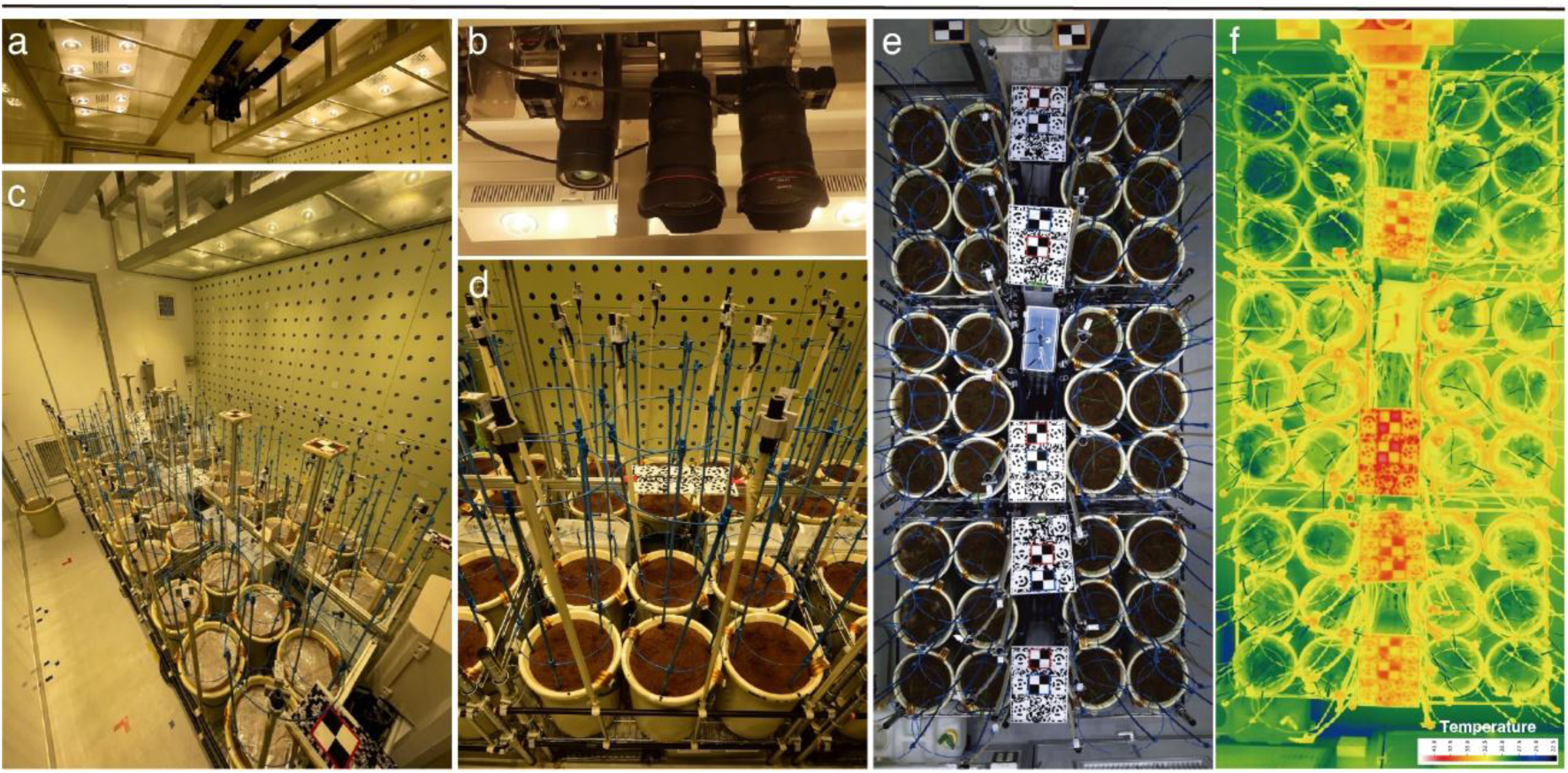
Overview of iPUPIL. **a.** A multi-camera system was installed on the ceiling. The multi-camera system moves forward and backward along a motorized linear stage. **b.** The multi-camera system comprises RGB, thermal, and monochrome cameras. **c.** Overview of the inside of iPUPIL. Thirty-six plant pots can be evaluated simultaneously. **d.** Front view of one unit of the automatic irrigation system. **e,f**. The entire RGB (**e**) and thermal (**f**) images of iPUPIL were created by combining photo series using the multi-camera system

A multi-camera system with up to four cameras was installed on top of each growth chamber to capture a series of plant images throughout the chamber (Fig. 1a,b). RGB and thermal cameras were installed to monitor changes in shoot growth and leaf temperature, indicators of drought tolerance^29^, respectively. The multi-camera system can be moved using a motorized linear stage placed on the ceiling. By repeatedly capturing images and moving, the system automatically acquires full plant images in the growth chamber according to a specific program (Fig. 1e,f, Supplementary Fig. 2). To evaluate shoot growth until harvest over time in long-growing crops such as rice, we modified the automatic irrigation system of the iPOTs to be more tolerant of long-term cultivation than its original mode (see Methods; Supplementary Methods, Supplementary Fig. 3). The environmental sensor system recorded the conditions in each pot during cultivation (Supplementary Fig. 4).

### Characterization of drought stress responses in cultivated and wild rice

We characterized the drought responses of five wild and four cultivated rice using iPUPIL (Table 1; see Methods). We conducted a 14-day drought treatment by stopping irrigation from 28 to 42 days after sowing (DAS). This period corresponded to the vegetative growth stage for all rice accessions (Fig. 2a, Supplementary Fig. 5). Compared with the soil water content (SWC) of the control plot, that of the drought plot decreased after irrigation was stopped but recovered after rewatering (Fig. 2a). Rice plants continued to grow well under well-watered conditions (WW). However, shoot growth was delayed and leaf temperature increased during drought treatment (DT) (Fig. 2b,c).

**Figure 2.**
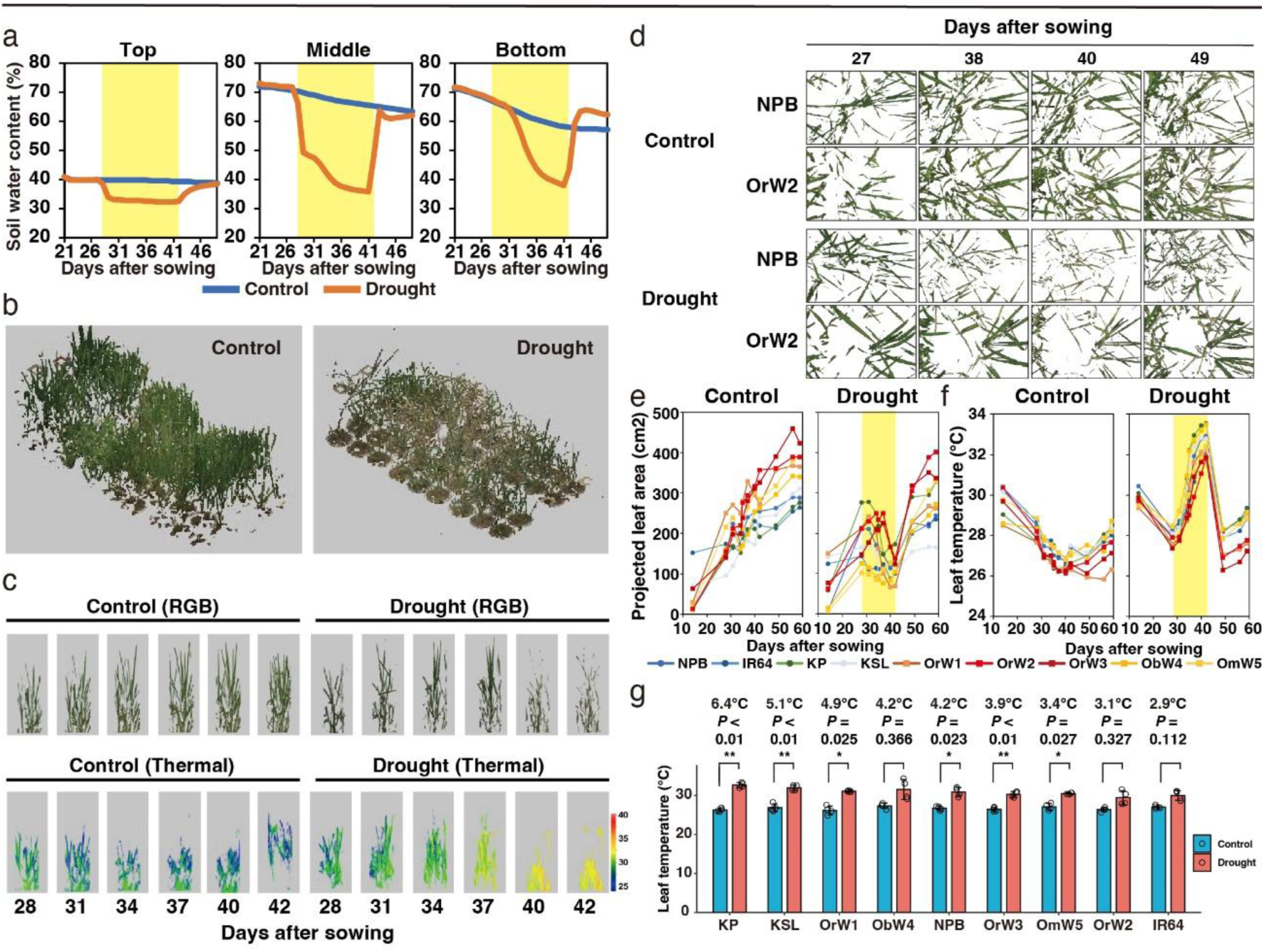
Morpho-physiological features of nine rice accessions under well-watered and drought conditions. **a.** Time course of mean soil water content at depths of 2 (top), 12 (middle), and 22 cm (bottom) from the soil surface in the control (*n* = 36) and drought (*n* = 36) plots. **b.** Reconstructed three-dimensional (3D) models 43 days after sowing (DAS) of plants grown in control and drought plots. **c.** Representative reconstructed 3D models of RGB from (upper) and thermal (lower) images of NPB. **d.** Representative images of reconstructed NPB and OrW2 plants in the control and drought plots. **e.** Time course of the projected leaf area (cm^2^) of representative plants in the control and drought plots. **f.** Time course of the mean leaf temperature (°C) of each plant (*n* = 3–4) in the control and drought plots. Data from the four corners were excluded owing to their low quality. **g.** Leaf temperatures of nine rice accessions at 36 DAS in the control and drought plots. The bars are arranged in order of temperature differences. Bars represent the mean (*n* = 4) ± standard deviation (SD). Bonferroni-corrected *P* value (two-sided Welch’s t-test) and the degrees of temperature change are shown.

**Table 1.**
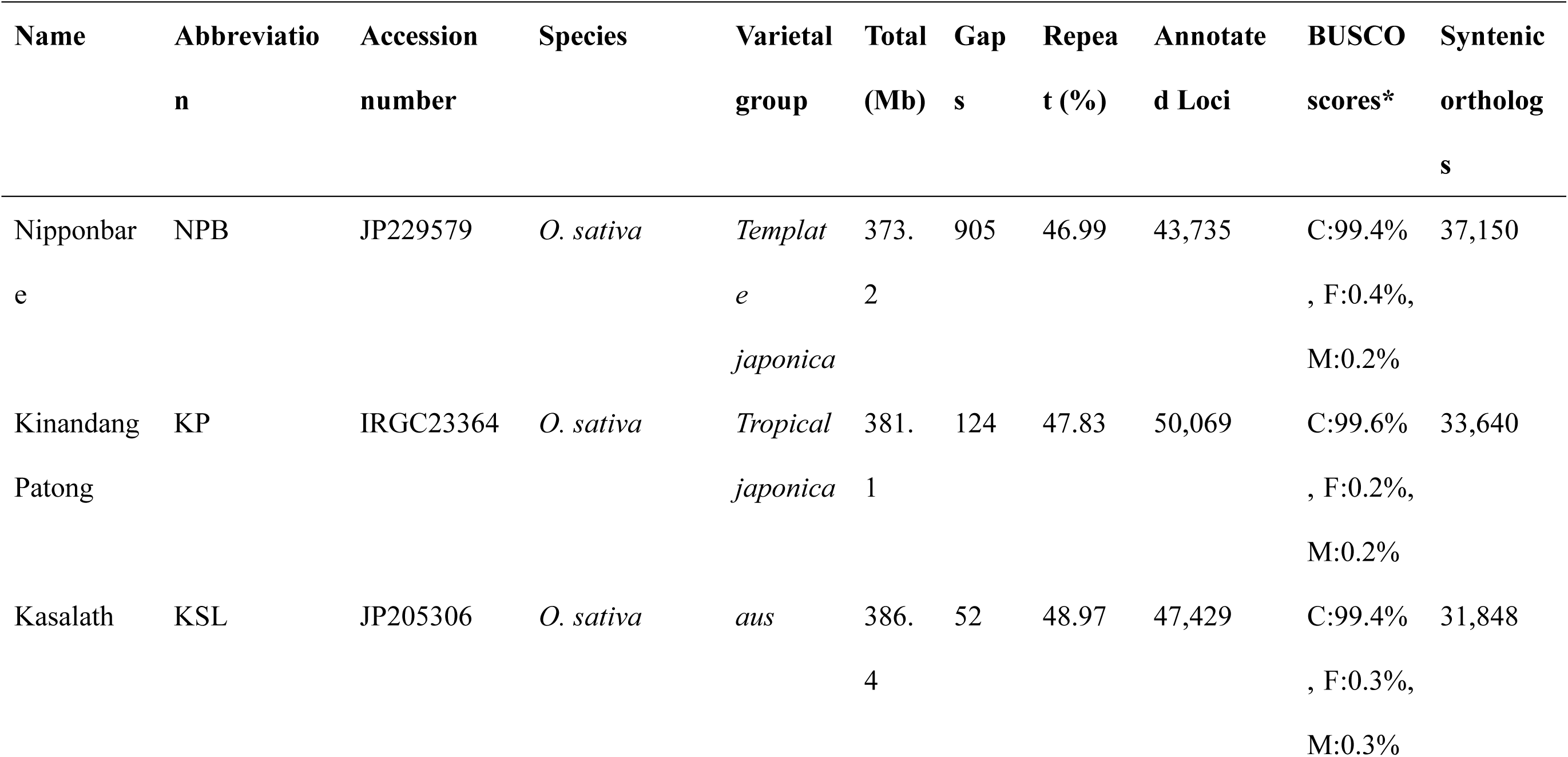
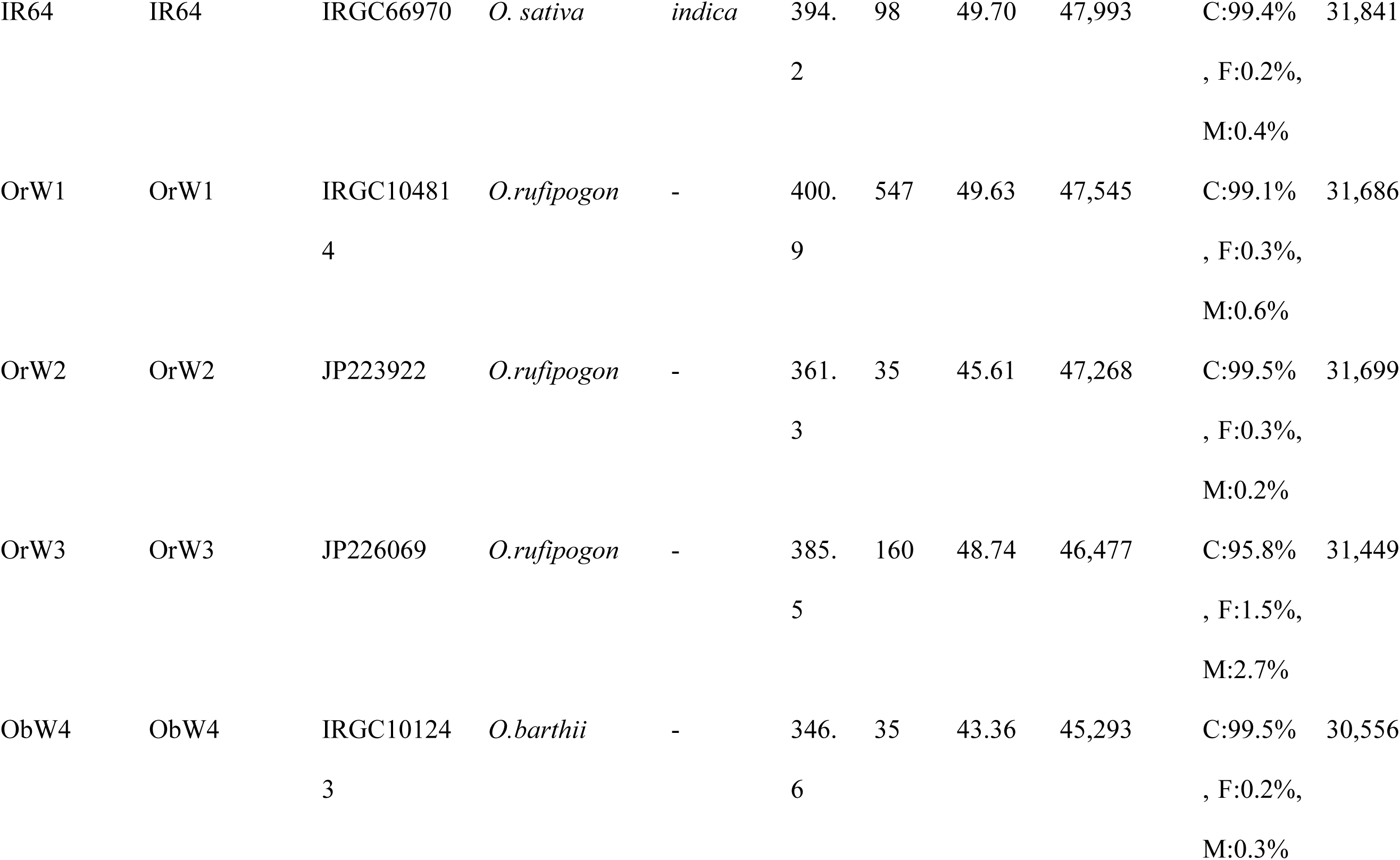
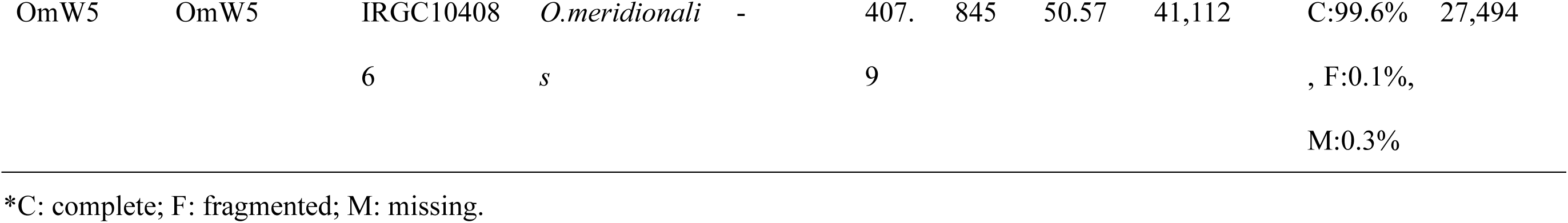
Summary statistics of assembly and annotation for nine Oryzeae reference genomes.

For morphological changes, we calculated the projected leaf area (PLA) based on RGB images (Fig. 2d, Supplementary Methods, Supplementary Fig. 6). In the drought plot, the PLAs of all accessions increased under WW. In contrast, PLAs decreased during DT and recovered after rewatering (Fig. 2e). The reduced PLA during DT was caused by growth retardation, including leaf rolling and wilting. The PLA of wild rice tended to be higher than that of cultivated rice under each condition (Fig. 2d,e). Among the wild rice accessions, OrW2 and OrW3 showed a small decrease in PLA during DT and increased PLA after rewatering (Fig. 2e). For example, OrW2 exhibited lower growth retardation under DT than the typical drought-sensitive cultivar, “Nipponbare” (NPB), as demonstrated by the RGB images and PLA (Fig. 2d,e).

We monitored leaf temperature to assess physiological changes (Fig. 2f). The leaf temperatures of all accessions tended to decrease in the control plot but increased during DT in the drought plot, indicating stomatal closure in response to drought (Fig. 2f). When comparing leaf temperature differences under DT relative to WW, “Kinandang Patong” (KP), “Kasalath” (KSL), OrW1, and ObW4 exhibited higher temperatures than those of NPB at 36 DAS, on the day with the greatest difference (Fig. 2g). In contrast, OrW3, OmW5, OrW2, and IR64 exhibited smaller temperature differences than those in NPB. Notably, OrW2 exhibited the lowest leaf temperature under DT (Fig. 2g).

### Time-course transcriptomic analysis identified wild rice with a unique drought response

In addition to NPB, which already has reference genomes, we constructed reference genomes and gene annotations for the eight rice accessions to analyze their gene expression profiles under drought conditions (see Supplementary Results, Supplementary Figs. 7–11). The evolutionary relationships of the nine accessions were inferred using the maximum likelihood method based on biallelic single-nucleotide polymorphism sites (Fig. 3a). These above-ground features varied widely, exhibiting erect to spread types (Fig. 3b).

**Figure 3.**
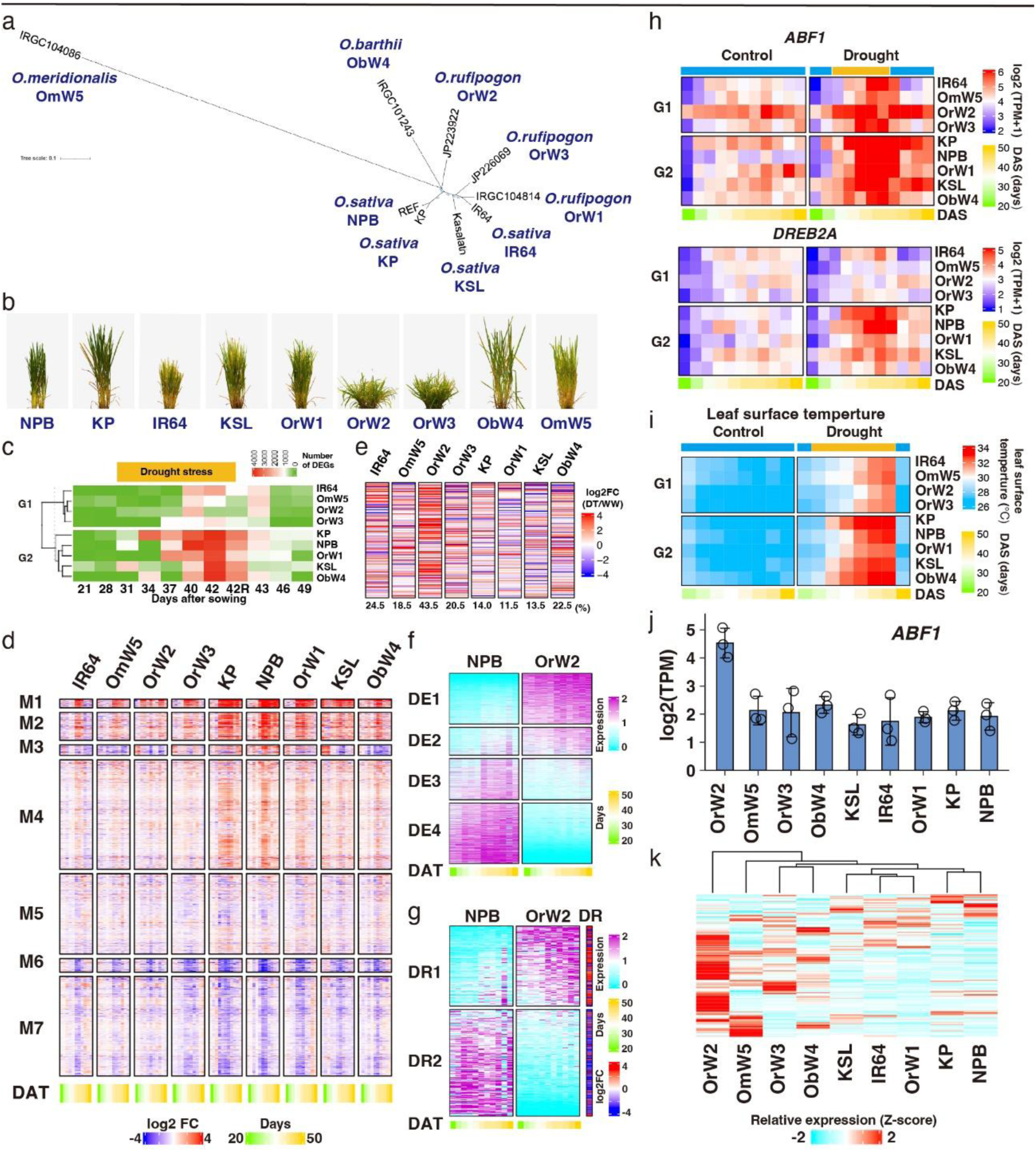
Genome and transcriptome features of nine rice accessions under control and drought conditions. **a.** Phylogenetic tree of nine rice accessions inferred using the maximum likelihood method. **b.** Images of rice varieties grown in iPUPIL. **c.** Heatmap of differentially expressed gene count (|log2FC| > 2, FDR < 0.05) in each rice plant in the drought plot compared with that in control plots. Rice accessions were classified into two groups: drought-insensitive (G1) and -sensitive (G2). **d.** Heatmap of relative drought stress-responsive gene expression levels in the drought plot compared with those in the control plot. Genes were classified into seven clusters according to drought stress responsiveness. **e**. Heatmap showing the drought stress responsiveness of 200 genes in NPB. In each accession, the top 200 genes upregulated before DT (21 DAS) compared with NPB were selected. The percentage under the heatmap indicates the proportion of drought-inducible genes. **f.** Heatmap of relative differentially expressed gene (|log2FC| > 2, FDR < 0.05) expression levels in OrW2 compared with those in NPB under control conditions. Genes were classified into four clusters according to the expression levels of OrW2 compared with those of NPB. **g**. Genes in genes listed in (**f**). Genes were classified into two clusters according to the expression levels of OrW2 compared with those of NPB. DR: relative expression level in NPB under control conditions compared with those under drought conditions. **h.** Heatmap of representative master regulator expression levels of drought stress-inducible genes, *ABF1* and *DREB2A,* in each rice accession under control and drought conditions. **i**. Heatmap of the projected leaf surface temperature, as shown in Fig. 2f. **j.** Expression levels of *ABF1* under WW. Bars represent mean (*n* = 3) ± standard deviation (SD). **k.** Heatmap of the relative expression levels of the downstream genes of ABF1 under WW. Z-score calculations were performed gene-wise. FC, fold change; FDR, false discovery rate; NPB, Nipponbare; KP, Kinandang Patong; KSL, Kasalath; WW, well-watered conditions; DT, drought treatment; DAS, days after sowing.

The differentially expressed gene (DEG) count in the drought plot compared with that in the control plot revealed that expression levels in OrW3, OmW5, OrW2, and IR64 were slightly altered. However, the expression of several genes in NPB, KP, KSL, OrW1, and ObW4 was remarkably altered (Fig. 3c). Notably, this drought response was similar to leaf temperature patterns during DT (Fig. 2g). Therefore, we classified the rice accessions into two groups: weakly (G1: OrW2, OrW3, OmW5, and IR64) and highly (G2: NPB, KP, KSL, OrW1, and ObW4) sensitive to drought (Fig. 3c). To clarify whether the expression patterns of drought-responsive genes differed between the two groups, we evaluated 3,145 drought-responsive genes that were one-to-one orthologous in nine accessions by classifying seven drought modules. Previously, drought-responsive genes in NPB have been classified into seven modules (M1–M7) based on their responsiveness to drought and functionality^12^. Drought-inducible genes involved in drought tolerance (M1, 2, and 4) and drought-repressible genes involved in plant growth, photosynthesis, and flowering (M6 and 7) did not show specific expression patterns in either accession (Fig. 3d). In contrast, these modules in G1 showed less expression variation under drought stress than those in G2.

The constitutive expression of specific genes before drought conditions affects the regulation of stress-responsive genes in cultivated rice^30^. We hypothesized that G1 accessions would exhibit constitutive expression of drought-responsive genes before DT, resulting in relatively small changes in gene expression after DT. To verify this hypothesis, we selected the top 200 genes that were more highly expressed in NPB before DT for each accession. The proportion of drought-inducible genes in NPB tended to be higher in G1 than in G2 (Fig. 3e). OrW2 exhibited a particularly high proportion of drought-inducible genes compared with that in the other G1 accessions. To elucidate the unique expression profile of OrW2, we compared the DEG expression levels in OrW2 to those in NPB in the control plot (Fig. 3f, Supplementary Fig. 12). The expression of genes related to photosynthesis, developmental processes, and multicellular organismal development was lower in OrW2 than in NPB (Fig. 3f). Next, we focused on DEGs with drought-responsive genes (Fig. 3g). The most highly expressed genes in OrW2 were drought-inducible and clustered within the DR1 module. In contrast, most downregulated genes in OrW2 were drought repressible and clustered within the DR2 module (Fig. 3g). Next, we assessed the expression levels of major transcription factors, which are master regulators of drought-responsive genes, including *ABF1/bZIP12*^9^, *DREB2A*^31^, and *NAC5*^32^. Their gene expression patterns resembled those of leaf temperature (Fig. 3h,i). Although the expression patterns of *DREB2A* and *NAC5* were similar in the nine accessions under WW, *ABF1* was constitutively highly expressed only in OrW2 (Fig. 3j, Supplementary Fig. 13). The expression pattern of *bZIP40*, the closest homolog of *ABF1* (Supplementary Fig. 14), was similar among the nine accessions (Supplementary Fig. 13).

Rice *ABF1* genes belong to subclass I, which is distinct from the *Arabidopsis* AREB/ABF group (subclass III), known regulators of ABA-mediated drought tolerance^33^. Rice ABF1 is a master transcription factor upstream of the subclass III AREB/ABF group, including OsbZIP23/OsAREB1/OsABF3, OsbZIP46/OsAREB8/OsABF2, and OsbZIP72/OsAREB2, which mainly regulate ABA-mediated drought-responsive genes^9,10,34^. ABF1-regulated transcripts^10^ accumulated in OrW2 under WW, whereas their levels remained low in other accessions (Fig. 3k). This suggested that constitutive *ABF1* expression was involved in the constitutive expression of drought-inducible genes in OrW2 plants under WW.

### Constitutive *ABF1* expression before drought is necessary for drought adaptation

We examined how constitutive *ABF1* expression helps OrW2 plants adapt to drought by comparison with NPB. After DT, both OrW2 and NPB plants exhibited reduced shoot lengths (Fig. 4a,b). OrW2 maintained its culm number, whereas that of NPB decreased (Fig. 4b). As a result, the shoot dry weight of OrW2 decreased by only 25%, compared with a 38% decrease in NPB (Fig. 4c). RT-qPCR analysis indicated that *ABF1* and the ABA-inducible gene *RAB16* were more constitutively expressed in OrW2 than in NPB, even under WW (Fig. 4d, Supplementary Fig. 15). The ABA content of OrW2 plants was similar to that of NBP plants in the control plot, suggesting that constitutive *ABF1* expression in OrW2 occurred in an ABA-independent manner (Fig. 4e). The drought survival assay showed that over 88% of OrW2 individuals survived severe drought conditions, whereas nearly all NPB plants almost died out (Fig. 4f). Constitutive *ABF1* expression may remain functional even under severe drought stress.

**Figure 4.**
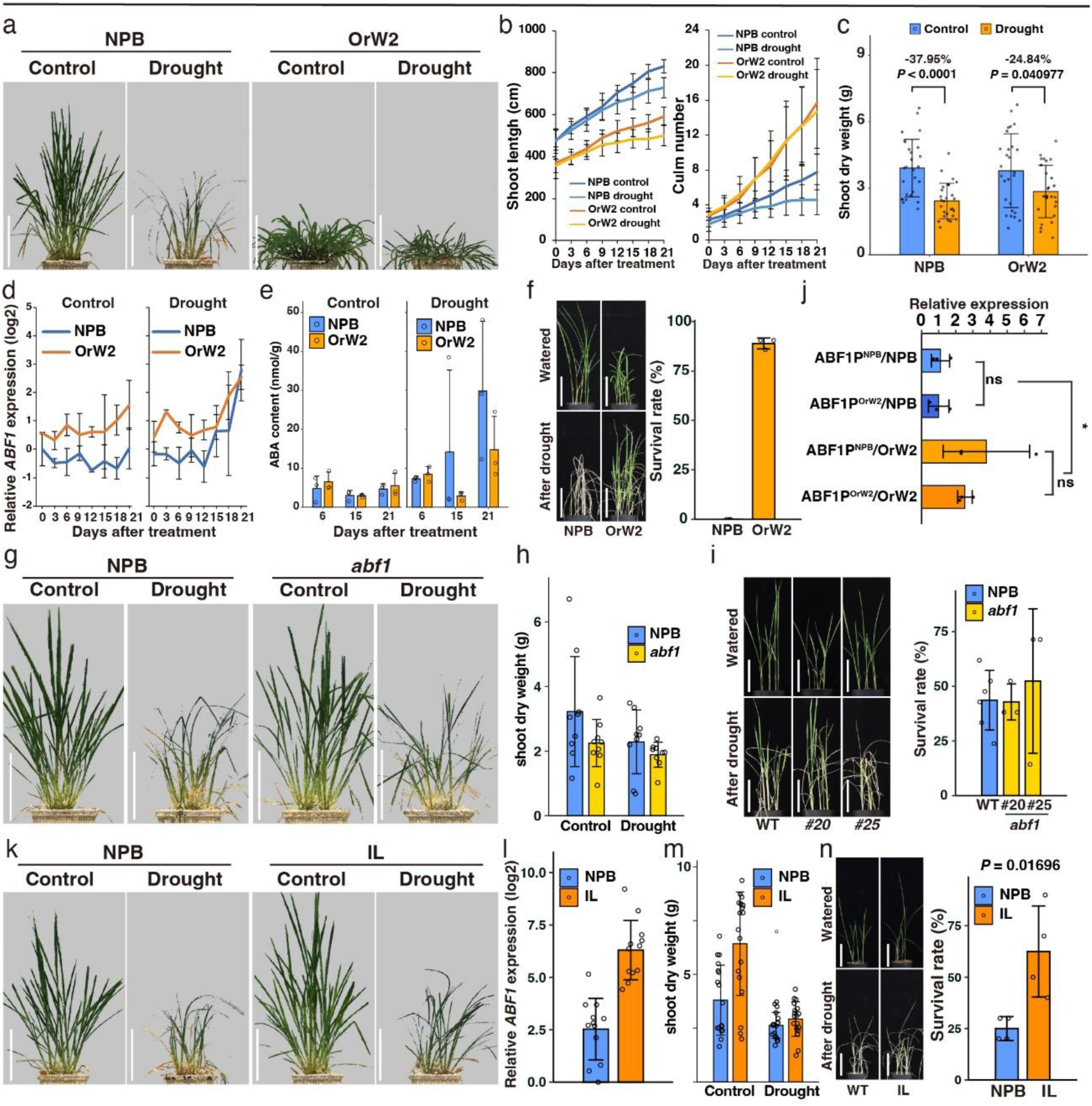
*ABF1* is constitutively expressed in drought-adapted rice. **a.** Representative growth of NPB and OrW2 in control and drought plots. Water supply stopped at 28 DAS (0 days after DT) until 49 DAS (21 days after DT). Scale bar (white) = 25 cm. **b.** Time course of shoot length and culm number per plant. Data represent the mean ± standard deviation (SD) (*n* = 27). **c.** Shoot dry weight of each rice plant grown in (**a**). Data represent the mean ± SD (*n* = 27). *P*-values are derived from a two-tailed Student’s *t*-test. Dry weight ratios of plants grown under drought/control are indicated. **d.** Time course of relative expression levels of *ABF1*. Data represent the mean ± SD (*n* = 3, biological replicates). **e.** ABA content of rice leaves in the control and drought plots. Bars represent the mean with ± SD (*n* = 3 biological replicates). **f.** Drought survival assay in NPB and OrW2. Data represent the mean ± SD (*n* = 3), based on three replicated experiments. Scale bar (white) = 10 cm. **g.** Representative growth of NPB (WT) and *abf1* in the control and drought plots. **h.** Shoot dry weight of WT and *abf1*. Data represent the mean ±SD (*n* = 9). **i.** Drought survival assay of *abf1.* Data represent the mean ± SD (*n* = 3), based on three replicates. **j.** The promoter assay was based on a transient expression test using protoplasts. Relative *ABF1* expression levels in protoplasts co-expressing *ABF1pro(NPB):ABF1* or *ABF1pro(OrW2):ABF1* are shown. Three biological replicates were used for each sample. Data represent the mean ± SD (two-tailed Student’s *t*-test: \**P* < 0.05). **k.** Representative growth of NPB and ILs in the control and drought plots. **l.** *ABF1* expression levels in NPB and ILs before DT (28 DAS). Data represent the mean ± SD (*n* = 12). **m.** Shoot dry weight of NPB and ILs grown in the control and drought plots. **n.** Drought survival assay of NPB and ILs. Data represent the mean ± SD (*n* = 3), based on three replicates (two-tailed Student’s *t*-test:). ABA, abscisic acid; NPB, Nipponbare; ILs, introgression lines; WT, wild-type; DT, drought treatment; DAS, days after sowing.

We evaluated the drought responses of *abf1* mutants developed using the CRISPR-Cas9 method (Supplementary Fig. 16, Fig. 4g). *abf1* mutant shoot length was similar to that of the wild-type (WT) in the control plot. In contrast, the culm number of *abf1* mutants was lower than that of WT (Supplementary Fig. 17). Shoot dry weight of *abf1* mutants decreased in the control and drought plots, suggesting the importance of *ABF1* in shoot growth, regardless of soil water conditions (Fig. 4h). However, drought survival assays showed no difference in survival rates between WT and *abf1* mutants (Fig. 4i). Under DT, where approximately half of NPB individuals die, the loss of *ABF1* function may not influence drought tolerance. This suggests that constant *ABF1* expression under WW, i.e., before DT, plays an important role in drought adaptation in OrW2.

### Dissection of constitutive *ABF1* expression in OrW2

To elucidate the reason for constitutive *ABF1* expression in OrW2, we examined the mutations and structural variants surrounding and within *ABF1*. Structural variations were observed only at points far from *ABF1* in OrW2 (Supplementary Fig. 18a). Selective sweep analysis revealed a signature in *ABF1* in *japonica* based on the composite likelihood ratio but not the fixation index, indicating that *ABF1* was not involved in artificial selection during domestication (Supplementary Fig. 19). In addition, the ABF1 protein sequences of NPB and OrW2 did not differ (Supplementary Fig. 20). In contrast, we identified a duplication of the upstream sequence of *ABF1* only in OrW2 (Supplementary Fig. 18b). Moreover, the upstream sequences of *ABF1* in OrW2 were distinct from those in *O. sativa* and other *O. rufipogon* accessions (Supplementary Fig. 18c). To clarify whether sequence differences in the promoter regions between OrW2 and NPB were involved in regulating constitutive *ABF1* expression, we selected two types of progenies from the BC_2_F_2_ introgression lines (ILs) derived from crossing NPB and OrW2: the NPB (ABF1p^NPB^) and OrW2 (ABF1p^OrW2^) homozygous alleles of the *ABF1* promoter region. *ABF1* expression between the ABF1p^OrW2^ and ABF1p^NPB^ lines under WW did not significantly differ (Supplementary Fig. 18d). Promoter assays of *ABF1* also showed no differences in the activities of ABF1p^OrW2^ and ABF1p^NPB^ in the NPB background (Fig. 4j). However, *ABF1* expression in the OrW2 background was higher than that in the NPB background, regardless of promoter type. These findings suggest the presence of an unknown trans-regulatory element that positively regulates *ABF1* expression in OrW2.

We obtained BC_2_F_3_ progeny from a BC_2_F_2_ line exhibiting constitutive *ABF1* expression, which was assumed to harbor both *ABF1* and unknown genes expected to regulate *ABF1* expression. We assayed the response to mild drought in BC_2_F_3_ ILs (Fig. 4k). *ABF1* and *RAB16* in BC_2_F_3_ ILs were constitutively expressed in WW (Fig. 4l, Supplementary Fig. 21). BC_2_F_3_ ILs showed superior growth than that of NPB in the control plot, although no significant difference in shoot biomass was observed between NPB and BC_2_F_3_ ILs in the drought plot (Fig. 4m, Supplementary Fig. 22). However, the drought survival assay revealed that BC_2_F_3_ ILs had higher survival rates than did NPB under severe drought conditions (Fig. 4n). Our findings suggest that the drought adaptability of OrW2 results from constitutive *ABF1* expression, facilitated by its interaction with unidentified factors that regulate *ABF1*.

## Discussion

Plant growth is typically repressed under drought conditions because of the tradeoff between growth maintenance and drought tolerance^35^. By suppressing growth, plants can conserve water and energy under drought conditions, thereby enhancing their survivability^36^. However, excessive growth inhibition in response to drought can lead to reduced crop yields. For example, drought tolerance has been improved by overexpressing drought-tolerant genes, such as *ABF1* and *DREB1*; however, most of these genes result in growth retardation due to decreased gibberellin (GA) content^37–39^. In contrast, in the current study, constitutive *ABF1* expression, without extreme expression levels in OrW2 and NPB backgrounds, improved drought tolerance while maintaining plant growth under normal conditions. Optimal and consistent *ABF1* expression may mitigate the negative relationship between shoot growth performance and drought resistance, resulting in drought tolerance without growth penalties in well-watered environments.

For the first time, plants exposed to stress were able to maintain pre-activated or sustained stress-related gene expression, allowing them to respond more quickly and efficiently than they would under typical stress responses, which are triggered only after stress occurs^40^. This priming-induced acquired tolerance likely provides wild plants with a significant adaptive advantage, enabling them to respond rapidly to unpredictable environmental fluctuations. However, our results do not support the interpretation of constitutive *ABF1* expression in OrW2 as a priming effect. In the control plot, OrW2 was unlikely to have experienced drought-induced priming because the automated irrigation system maintained well-watered conditions. Furthermore, analysis of *ABF1* expression in ABF1p^OrW2^ and ABF1p^NPB^ ILs (Supplementary Fig. 18d) suggested that epigenetic regulation was not responsible for constitutive *ABF1* expression in OrW2. Thus, the regulatory mechanism of *ABF1* expression identified in OrW2 is poorly understood. However, one possibility is the existence of a trans-element that regulates *ABF1* expression. A gain-of-function mutant of the DELLA protein SLENDER RICE 1 (SLR1), a negative regulator of GA signaling, exhibited enhanced drought tolerance associated with low *ABF1* expression levels^36^. Notably, a loss-of-function mutant of *OsJAZ6*, a negative regulator of tillering and drought responses that interacts with SLR1 to promote its degradation, displayed enhanced drought tolerance and increased tiller number, similar to OrW2^41^. These suggest that the SLR1-related pathway is partially involved in *ABF1* expression in OrW2. Further studies using single-cell analysis and/or hormone profiling are required to verify this hypothesis. Another possibility is that the upstream osmotic sensing mechanisms controlling *ABF1* expression are enhanced in OrW2. The RAF-SnRK2 pathways^42–48^, central pathways in ABA-dependent or -independent drought stress signaling, may be constantly active in OrW2. A detailed activation assay for these kinases may contribute to elucidating the molecular mechanisms underlying constitutive *ABF1* expression in OrW2.

A stable drought response is a key adaptation for wild species inhabiting disturbed areas with recurring dry–wet cycles. For instance, OrW2, collected during our wild rice exploration in Myanmar^49^, naturally grows in disturbed swamps with alternating dry and wet conditions, adjacent to rice paddies (Supplementary Fig. 23). During early-dry season swamp visits, we observed that swamps near paddy fields remained damp, whereas those farther away were extremely dry. Given the significant soil moisture variability in its habitats owing to changing climate and weather, this adaptive mechanism, involving constitutive *ABF1* expression, likely evolved to help OrW2 cope with persistent drought stress. Phylogenetic analysis indicated that OrW2 differentiated earlier than the other *O. rufipogon* accessions, such as OrW1 and OrW3 (Supplementary Fig. 10). Therefore, we speculated that a unique adaptation strategy to drought stress is likely to occur in OrW2, leveraging drought-tolerance genes, including *ABF1*.

A comparative analysis of the phenotypic and transcriptomic data obtained in iPUPIL revealed that constitutive *ABF1* expression, found only in OrW2, is an adaptive mechanism that may contribute to the ability of wild rice to adapt to disturbed areas under repeated dry–wet conditions. Understanding these regulatory mechanisms will contribute to developing drought-resistant rice that does not incur yield penalties. In addition, our unmanned phenotyping platform offers a new approach for screening plants with unique drought stress responses. A comprehensive genomic dataset, including time-course transcriptomic and phenotypic data of wild rice accessions, is expected to provide insights into the undiscovered mechanisms of drought tolerance, ultimately informing the development of drought-resistant rice.

## Online Methods

### iPUPIL configuration

We constructed two custom-designed growth chambers that could be controlled independently (Nippon Medical & Chemical Instruments Co., Osaka, Japan). The internal dimensions of the growth chambers were 2,118 mm in width, 2,850 mm in depth, and 2,000 mm in height. Environmental parameters could be controlled separately in each chamber, including air temperature (20–45°C) and humidity (30–50%). Multi-camera, automatic irrigation, and environmental sensor systems for each plant pot were installed inside the chamber. Each growth chamber accommodated the evaluation of 36 plant pots. The multi-camera system could accommodate up to four different camera types.

In this study, RGB (LUCIS Atlas 10; LUCID Vision Labs Inc., Burnaby, BC, Canada) and infrared (FLIR A615; Teledyne FLIR LLC., Wilsonville, OR, USA) cameras were used. This multi-camera system was mounted on a motorized linear stage fixed to the ceiling, allowing movement from the start (0 mm) to the end (2,240 mm) of the growth chamber. Movement and photography were performed automatically using a specific program (Supplementary Fig. 2).

The environmental sensor system comprised aerial and soil sensors. The arial sensor, positioned atop each pot, recorded humidity, temperature, and light intensity, while two soil sensors were installed per pot. These sensors could detect soil water levels at depths of 2 cm (top), 12 cm (middle), and 22 cm (bottom) from the soil surface. SWC was calculated as previously described^25^.

### Plant materials

We used four cultivated rice accessions (*O. sativa* L.), Nipponbare (temperate *japonica*), Kinandang Patong (IRG23364; tropical *japonica*)^49^, IR64 (IRGC66970; *indica*)^49^, and Kasalath (*aus*), and five wild rice accessions, *O. rufipogon* Griff. (IRGC104814, JP223922, and JP226069)^16,50,51^, *O. barthii* A.Chev. (IRGC101243), and *O. meridionaris* Ng (IRGC104086) (Table 1).

### Mild drought test in iPUPIL

We evaluated the drought responses of the nine rice accessions using iPUPIL. Rice seedlings were grown in growth chambers under a 14 h light/10 h dark photoperiod with 50% relative humidity. Light intensity gradually changed during this stage (Supplementary Fig. 4). The temperature in the growth chambers varied from 25°C (night) to 30℃ (daytime) as previously described (Supplementary Fig. 4). Calcined clay (Profile® Greens GradeTM; PROFILE Products LLC, Buffalo, IL, USA) was used instead of soil media. The calcined clay was rinsed with water and dried before use. After filling the pot with 9 L of calcined clay, we supplied it with a modified Kimura B hydroponic solution before sowing as previously described^12^.

In the control plots, the water level in each pot was maintained at 10 cm from the bottom until 14 DAS, after which height was adjusted to 8 cm, representing well-watered (WW) conditions. In the drought plot, rice plants were grown under the same irrigation conditions as those in the control plot until 28 DAS. Water supply was stopped at 28 DAS to implement DT, and rewatering occurred at 42 DAS. The DT conditions were determined based on variation in initial growth rates and phase change timing of vegetative growth-reproductive growth among rice accessions. We applied DT during the 2-week period from 28 to 42 DAS, which is considered the vegetative growth stage for all rice accessions (Supplementary Fig. 5). In a previous study, we found that drought stress during this growth period significantly affected rice growth^12^.

### Sample and library preparation for RNA-seq

The youngest, fully expanded leaves of each rice accession grown in iPUPIL were collected for RNA extraction. Total RNA was extracted from shoot samples using guanidinium thiocyanate and isopropyl alcohol-based RNA extraction methods, as previously described^52^. Equal amounts of RNA extracted from three independent plants in the same pot were used for RNA-seq library preparation, which was conducted using a NEBNext Ultra II Directional mRNA-seq kit (New England Biolabs, Ipswich, MA, USA) according to the manufacturer’s protocols as previously described^12^. These libraries were sequenced using an Illumina NovaSeq 6000 (Illumina, San Diego, CA, USA) at Macrogen (Seoul, South Korea). Sequencing was performed in S4 flow cells with 150-bp paired-ends and unique dual-index reads, as previously described^12^.

### RNA-seq data analysis

Paired-end RNA-seq reads were aligned to the corresponding genome sequences using HISAT2 with the aforementioned parameters. Gene abundance (TPM values) for each gene and transcript was estimated using StringTie (v2.2.1)^53^. Heatmaps of gene expression levels (Z-score-normalized TPM) were visualized using the R package ComplexHeatmap^54^. Candidate genes directly regulated by OsABF1 were identified from a previous study^10^.

To elucidate transcriptomic insights into drought response in the nine accessions, we collected 594 leaf samples (11 time points, 2 environmental conditions, and 9 rice accessions, with 3 biological replicates) for RNA-seq analyses. Four samples (Control 27 DAS KP, Drought 34 DAS OrW1, Drought 34DAS OrW2, Control-Rewatering 43 DAS OmW5) were excluded from analysis because of low library quality.

## Acknowledgments

We thank S. Yamazaki, H. Minowa, K. Okumura, M. Heta, T. Nagashima, and T. Shinotsuka (Tecs Inc., https://www.tecs.ne.jp) for supporting the setup of the multi-camera system; M. Matsuyama (ViewPLUS Inc., https://www.viewplus.co.jp/) for providing technical support to develop the program of camera control; I. Kijihana, T. Nishi, Y. Kanaya, and N. Sasaki (NIPPON MEDICAL & CHEMICAL INSTRUMENTS CO., LTD., https://www.nihonika.co.jp/) for supporting the setup of a growth chamber; M. Endo (NIAS, NARO) for providing the vectors used for genome editing; our staff (N. Kanno, Y. Fukuda, M. Iba, H. Tanaka, C. Kawashima, K. Onoe, and S. Yabe) for their technical assistance in plant cultivation and phenotyping. This work was supported by the Cabinet Office, Government of Japan; the Moonshot Research and Development Program for Agriculture, Forestry, and Fisheries (funding agency: Bio-oriented Technology Research Advancement Institution, grant number JPJ009237); and JST CREST (grant number JPMJCR17O1).

