## Supplementary Information for "Multi-omics profiling with indoor-unmanned phenotyping reveals drought adaptation through constitutive *ABF1* expression in wild rice"

**Contents**

Supplementary Results

Supplementary Methods

Supplementary Figures 1-23

Supplementary References

#### Supplementary Results

##### *De novo* assembly and gene annotation in cultivated and wild rice genomes

The HiFi reads were assembled *de novo* into contig sequences, which were scaffolded using optical mapping data to obtain chromosome-scale genome sequences. Each assembled genome sequence demonstrated a level of contiguity and completeness comparable to or higher than that of the Nipponbare (NPB) reference genome (IRGSP-1.0) (Table 1). A consensus repeat library was constructed based on the reference genome of NPB and the assembled genomes of eight rice accessions, and the repeat content of each genome was investigated. The proportion of repetitive sequences in the genomes of the eight rice accessions ranged from 43.4 to 51.0%, showing a clear positive correlation with genome size (Pearson's correlation coefficient = 0.996, $P < 0.001$) (Table 1). Retroelement-derived repeats were particularly abundant among the repetitive sequences. Notably, the number of Gypsy-type retrotransposons was positively correlated with genome size (Pearson's correlation coefficient = 0.915, $P < 0.001$).

The genome structures of NPB and eight other rice accessions were similar (Supplementary Fig. 7). A relatively large difference observed in the plot was an inversion near the centromeric region of chromosome 6. OmW5 also showed a relatively large inversion on chromosome 4. To further investigate the diversity of these genome sequences, a pangenome graph was constructed for the nine rice accessions. Using the NPB genome as a reference, the single-nucleotide polymorphisms (SNPs) and structural variants (SVs) in each genome sequence were analyzed. A large number of relatively large insertions and deletions that were difficult to detect using short-read-based analysis were identified (Supplementary Fig. 8 and 9).

*Ab initio* gene prediction was performed using RNA-seq and known protein sequences. NPB reference gene models were mapped to the genome. The predicted gene models were integrated with the reference gene models to annotate the eight reference genomes. The number of predicted loci in each genome ranged from 41,112 to 50,069. As reported by Stein et al.[1], the phylogenetic tree indicated that *Oryza sativa* and *Oryza rufipogon* formed a cluster, while ObW4 and OmW5 were located outside this cluster, with OmW5 being the most distantly related (Supplementary Fig. 10). Based on gene clustering based on amino acid sequence similarity, 51,038 orthogroups were identified in the 9 accessions. Of these, 24,258 (47.6%) orthogroups were conserved in all nine accessions (Supplementary Fig. 11). In contrast, single-copy orphan genes, which were present in 1 accession, were also found in other accessions (2,218, 737, 445, 415, 429, 814, 493, 1,090, and 1,727 genes for NPB, Kinandang Patong (KP), Kasalath (KSL), IR64, OrW1, OrW2, OrW3, ObW4, and OmW5, respectively). The one-to-one gene relationships between varieties were estimated using synteny information. As a result, a total of 37,281 genes were identified as having one-to-one correspondence across the 9 accessions. To enhance interpretability, these one-to-one orthologous genes were used in transcriptome analysis of the nine accessions. Chromosome-scale reference genomes, gene annotations, and ortholog information can be valuable resources for exploring genes involved in various stress tolerance mechanisms.

#### Supplementary Methods

##### Modification of the bottom-irrigation system iPOTs

We used an automatic irrigation system (Tecs Inc., Ibaraki, Japan), which is an improved version of iPOTs[2]. Six pots in each unit were watered using a uniform adjustment value, achieved by lowering the center of gravity per unit. Water levels were automatically controlled using a specific program. The influence of water management on the plants was minimized by installing a water supply and drainage system outside the chamber. We also modified the environmental sensor system to measure the above- and underground environmental values for each pot (see Methods; Supplementary Fig. 3). The aboveground sensors monitoring temperature, humidity, and light intensity were installed at a height of 1,400 mm from the top of the pot (Supplementary Fig. 1). To measure soil water content, two sensor sheets were attached to the inner surface of the pot (Supplementary Fig. 3).

##### 3D reconstruction of aboveground architecture

To estimate the leaf area and leaf surface temperature of individual plants during cultivation, 3D point clouds were created using images captured automatically using a multi-camera system. A total of 113 RGB and 113 thermal images were acquired, and 3D point clouds for the measurements were constructed using 3D reconstruction and registration processes (Fig. 2b, Supplementary Fig. 6). The leaf area was estimated from an RGB image projected from an overhead viewpoint (Supplementary Fig. 6e). Leaf surface temperature was estimated as the average temperature of the points (Supplementary Fig. 6f). The 3D-reconstructed leaf area was estimated using only representative plant pots, to prevent the inclusion of pots at the edges of rooms with insufficient reconstruction (Fig. 2e). To exclude non-plant temperatures sources, the leaf temperature of plants enclosed within the cylinder was estimated (Fig. 2f, Supplementary Fig. 6).

In the 3D reconstruction process for the aboveground architecture of rice plants, 3D point clouds were reconstructed using Structure from Motion[3] and Multi-View Stereo[4] techniques, based on RGB and thermal imagery. During the registration process, point clouds reconstructed from RGB (RGB point cloud) and thermal (thermal point cloud) images were registered at the same spatial location.

During the measurement process, point clouds were extracted per plant pot, and the leaf area and temperature were measured. The projected leaf area was measured from an image in which the RGB point cloud was extracted as a cuboid region and projected onto an overhead viewpoint. Leaf temperature was measured as the average temperature of a thermal point cloud, which was extracted as a cylindrical region (16 cm in diameter). These processes are conducted in Python (3.8). The Metashape Python module (Agisoft LLC, "Metashape Professional 1.8.5", http://www.agisoft.com/, (Accessed Feb 3, 2025)) was used for Structure from Motion, Multi-View Stereo and registration. Plant 3D point clouds were visualized using CloudCompare (CloudCompare 2.13.0; https://www.cloudcompare.org/)[5]. All-around 3D plant modeling using multiple images was conducted using an improved Structure from the Motion/Multi-view Stereo method[6].

**Sample and library preparation for genome sequencing**

Total DNA was extracted from the leaves of each rice accession grown in greenhouse under natural day condition using a DNeasy Plant Mini Kit (Qiagen, Hilden, Germany).

DNA libraries were prepared using a SMRTbell Express Template Prep kit 2.0 (Pacific Biosciences, Menlo Park, CA, USA) and sequenced in the HiFi mode of the PacBio_Sequel II platform (Pacific Biosciences). For optical mapping, DNA samples extracted using Plant DNA Isolation kit (Takara Bio, Shiga, Japan) and were labeled using a DLS DNA Labeling Kit (Bionano Genomics, Inc., San Diego, CA, USA) and subsequently loaded onto a Saphyr system (Bionano Genomics, Inc.).

#### ***De novo* assembly**

For each variety, HiFi reads generated by PacBio sequencing were assembled into contigs using Hifiasm (v0.16.1)[7]. Redundant haplotypes were identified and removed using the purge_dups pipeline (v1.2.5, https://github.com/dfguan/purge_dups) with minimap2 (v2.24)[8]. Homology-based misassembly correction was performed using RagTag (v2.1.0)[9,10] based on the reference genome of NPB (IRGSP-1.0) and HiFi reads. Hybrid scaffolding was performed with both the contigs and Bionano optical mapping data using the Bionano Solve pipeline (v1.0). First, the contig sequences were converted into a CMAP file through *in silico* digestion with the DLE-1 enzyme, and the HybridScaffold script was executed with the parameters -B 2 -N 2. Scaffold sequences were aligned and anchored to each chromosome using RagTag, with the IRGSP-1.0 genome as the reference. Organellar genome-derived contigs were removed by mapping them to the mitochondrial and chloroplast genomes using minimap2 under the following criteria: alignment was primary alignment, mapping quality exceeded 20, more than 90% of the contig sequence was aligned, and the mismatch rate of the alignment was below 0.1%.

**Gene annotation**

In the singularity environment, repeat masking of the reference genomes was conducted using the Dfam TE Tools Container v1.6. The consensus repeat library was constructed using RepeatModeler (v2.0.4) based on the IRGSP-1.0 reference genome and eight assembled genome sequences. Repetitive sequences in the eight genomes were masked using RepeatMasker (v4.1.3). For gene prediction, the known protein sequences of *Oryza* species were downloaded from UniProt (https://www.uniprot.org/). RNA-seq reads were aligned to each genome using HISAT2 (v2.2.1) with the following parameters: --min-intronlen 20 –max-intronlen 10000[11]. Protein-coding genes were predicted using BRAKER3 (v.3.0.3) with known protein sequences and RNA-seq read alignment[12]. The gene annotation of the IRGSP-1.0 genome was downloaded from RAP-DB (https://rapdb.dna.affrc.go.jp) and RGAP (https://rice.uga.edu) databases and used as a reference. Reference gene models were mapped to eight genomes using Liftoff[13]. Two gene model sets predicted using BRAKER3 and Liftoff were merged, and redundant gene models were removed using gffreads with M-K-Q options and in-house scripts[14]. Functional domains were predicted for each protein sequence, Gene Ontology (GO) terms were assigned using InterProScan (v5.61-93.0), and homologous known protein sequences were searched against the UniProt database using DIAMOND (v. 2.1.8)[15,16]. BlastKOALA was used to assign KEGG Orthology terms to each protein sequence[17]. For each locus, the transcript with the most functional annotations and longest amino acid sequence was selected as the primary transcript. The completeness of the assembled genome and gene models was evaluated using BUSCO (v5.7.1) with the Embryophyte _odb10 dataset.

**Comparative gene and genome analysis**

Pangenome graphs and whole-genome alignments of the nine genomes were constructed using Minigraph-Cactus (v2.9.2)[18]. Genome graph data were converted into variation data using the NPB genome as a reference, and SNPs and InDels in each genome were detected. A phylogenetic tree of the nine rice varieties was constructed using the maximum likelihood method in IQ-TREE. Genetic distances were estimated using 6,151,199 bi-allelic SNP sites. Syntenic orthologs conserved in a one-to-one relationship across the nine rice species were extracted using MCScanX in September 2023[19]. Orthogroups of the nine species were predicted using OrthoFinder (version 2.5.4)[20].

**Quantitative reverse transcription PCR (RT-qPCR) analysis**

cDNA was synthesized using a High-Capacity cDNA Reverse Transcription Kit (Thermo Fisher Scientific, Waltham, MA, USA) with random hexamer primers, according to the manufacturer's instructions. THUNDERBIRD® SYBR qRT Mix (TOYOBO, Osaka, Japan) was used for the quantitative RT-qPCR assay. The oligonucleotide primers used for *OsABF1*[21] and *OsRAB16A*[22] have been referred to in previous studies. Expression levels were calculated using the $2^{-\Delta\Delta CT}$ method[23].

**Mild drought assay in growth chambers.**

To analyze the drought tolerance mechanism of OrW2 in detail, we evaluated OrW2, NBP, introgression lines (ILs), and *abf1* mutants in a large growth chamber. The drought treatment (DT) was applied for 21 days from 28 DAS. Rice plants were grown under a 14 h light/10 h dark photoperiod. The temperature in the growth chambers was 30°C (daytime) and 25°C (night). The water supply was stopped at 28 DAS, and rewatering was conducted at 49 DAS. The shoot dry weight was measured after the samples were overdried at 80°C for 72 h as previously described[2].

**Measurement of abscisic acid (ABA) levels**

Expanded rice leaves were collected to measure ABA content. All samples were dried using a freeze dryer and crushed using beads. ABA was extracted using 80% methanol containing 500 mg/L citric acid. After centrifugation, the supernatants were collected and dried. ABA levels were measured using a Phytodetek ABA Measurement Kit (Agdia, Elkhart, IN, USA), according to the manufacturer's protocol.

**Drought survival assay**

To evaluate OrW2, NBP, and ILs, as well as the *abf1* mutants, we conducted drought survival assays using small pots[24]. Five to seven plants from each line were grown in a mixed soil comprising Bonsol No.1 (Sumitomo Chemical, Tokyo, Japan) and Ikubyo-Shibaue-Yodo (Shidara, Kanuma City, Japan). Next, 14 DAS plants were subjected to drought stress for 12–14 days, and the survival rate was calculated after rewatering for 7 days. Rice seedlings were grown in growth chambers under a 14 h light/10 h dark photoperiod with 50% relative humidity. The temperature in the growth chambers was 30°C (daytime) and 25°C (night).

***ABF1* mutant test using a CRISPR-Cas9 system**

For genome editing, CRISPR/Cas9 cleavage sites of *ABF1* were designed using CRISPR Direct[25], and the vector was constructed as previously described[26]. The construct was introduced into *Agrobacterium tumefaciens* strain EHA105 through electroporation, and *Agrobacterium*-mediated transformation of rice was performed as described previously[27]. Mutations in the genomic region of *ABF1* were identified using Sanger sequencing, and the homozygous mutants were designated *abf1*.

**Selective sweep analysis**

Genotype data for cultivated and wild rice accessions were obtained from the NARO Open Rice Collection (NRC)[28] and OryzaGenome release 2.1 (http://viewer.shigen.info/oryzagenome21detail/), respectively. Fixation index ($F_{ST}$) values were calculated using vcftools v.0.1.15[29,30] with a sliding window size of 100 kb, specified by the -fst-window size command, and a step of 10 kb, specified by the -fst-window step command. Considering the domestication process of cultivated rice, we calculated the $F_{ST}$ values between *indica* and Or-I and *japonica* and Or-III separately. The ecotype categorization of wild rice accessions (Or-I and Or-III) was based on Huang *et al.*[31]. Selective sweep analysis within domesticated rice accessions was performed based on the composite likelihood ratio (CLR) test using SWEED version 4.0.0, with default parameter settings[32]. In both $F_{ST}$ and CLR, the top 5% of the scores were used as the threshold for significance.

**Development of ABF1pOrW2-homo and ABF1pNPB-homo lines**

$F_1$ seeds were obtained by crossing NPB with OrW2. The ABF1pOrW2-homo and ABF1pNPB-homo lines ($BC_2F_2$) were developed by backcrossing $F_1$ plants with NPB twice, and the progenies were selected using PCR with primers to distinguish between ABF1pOrW2-homo and ABF1pNPB-homo.

**Protoplast assay**

The genomic region of *ABF1* was amplified using PCR with the NPB or OrW2 genome as a template and inserted into pCAMBIA1301 to generate pCAMBIA1301-ABF1p:ABF1 using an Infusion Directional Cloning Kit (Takara Bio, Shiga, Japan). For the protoplast assay, ABF1p:ABF1-NosT was amplified using PCR with pCAMBIA1301-ABF1 as the template, and the fragments were inserted into the pUC19 vector to generate pUC19-ABF1p:ABF1. Rice protoplasts were prepared from the stems and sheaths of 14 DAS seedlings[33] and hydroponic cultivation[34].

**Phylogenetic analysis**

Phylogenetic trees and multiple alignments were constructed from full-length protein sequences using MEGA11[35]. The phylogenetic tree was constructed using the neighbor-joining method[36].

**Supplementary Figures**

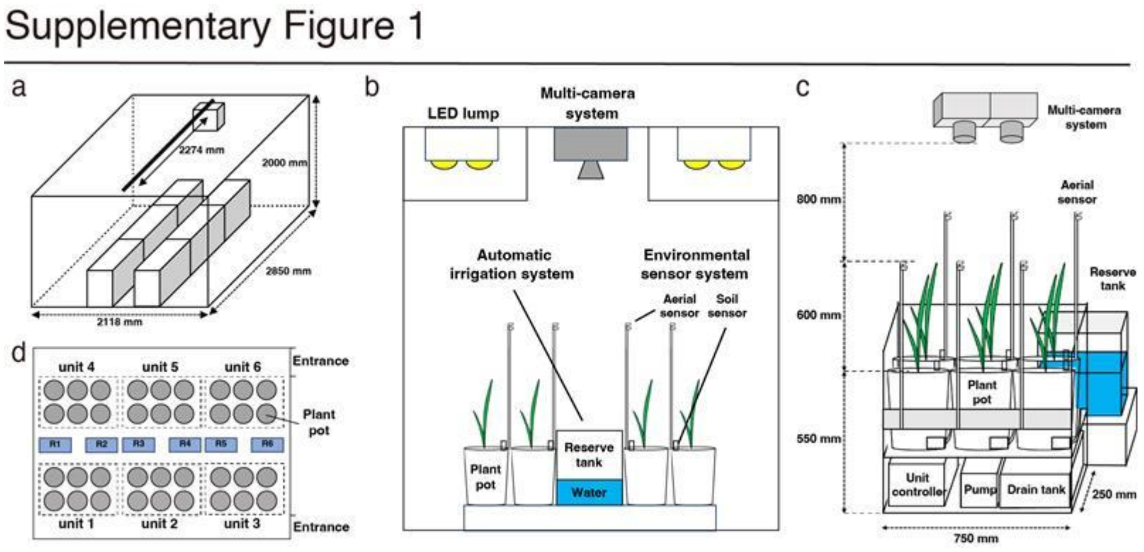


**Supplementary Figure 1. Schematic diagrams of iPUPIL.**

**a.** Simplified internal schematic of iPUPIL per growth chamber. The camera system moves along the rail indicated by the black line at the top of the chamber. The six boxes at the bottom represent individual irrigation systems. **b.** Front view of iPUPIL. The multi-camera system was installed between LED lights. The automatic irrigation system controls water level per unit, and an environmental sensor system for each pot was installed. The soil sensors were embedded in the pot soil. **c.** Side view of one unit of the automatic irrigation system in iPUPIL. Water stored in the reserve tank (shown in light blue) is automatically distributed to each pot. **d.** In the top view of the plant pots, the automatic irrigation system regulates the water level of six pots by connecting them to a single unit. Up to six units can be placed in iPUPIL. The gray circles represent individual plant pots, and the light blue boxes represent the reserve tanks.

#### Supplementary Figure 2

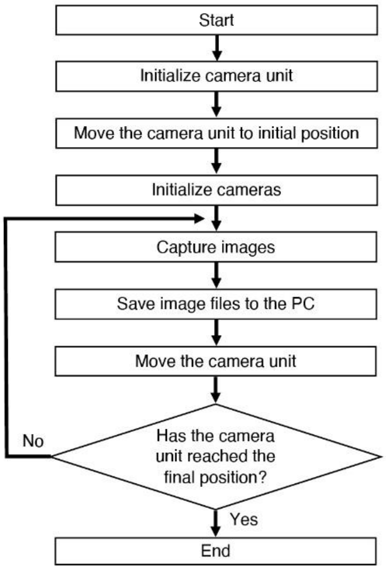


**Supplementary Figure 2. Image acquisition process of multi-camera system.**

A flowchart of the photography operation is presented. After initializing the device, image capturing and unit movement were repeated to acquire images.

#### Supplementaly Figure 3

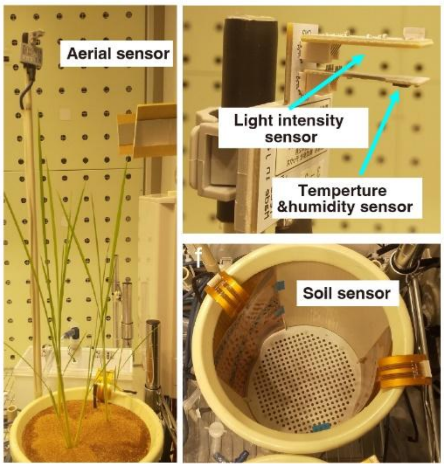


**Supplementary Figure 3. Environmental sensor system in iPUPIL.**

The aerial sensor comprises a light intensity sensor, as well as temperature and humidity sensors. Two soil-moisture sensors were placed in each pot.

#### Supplementary Figure 4

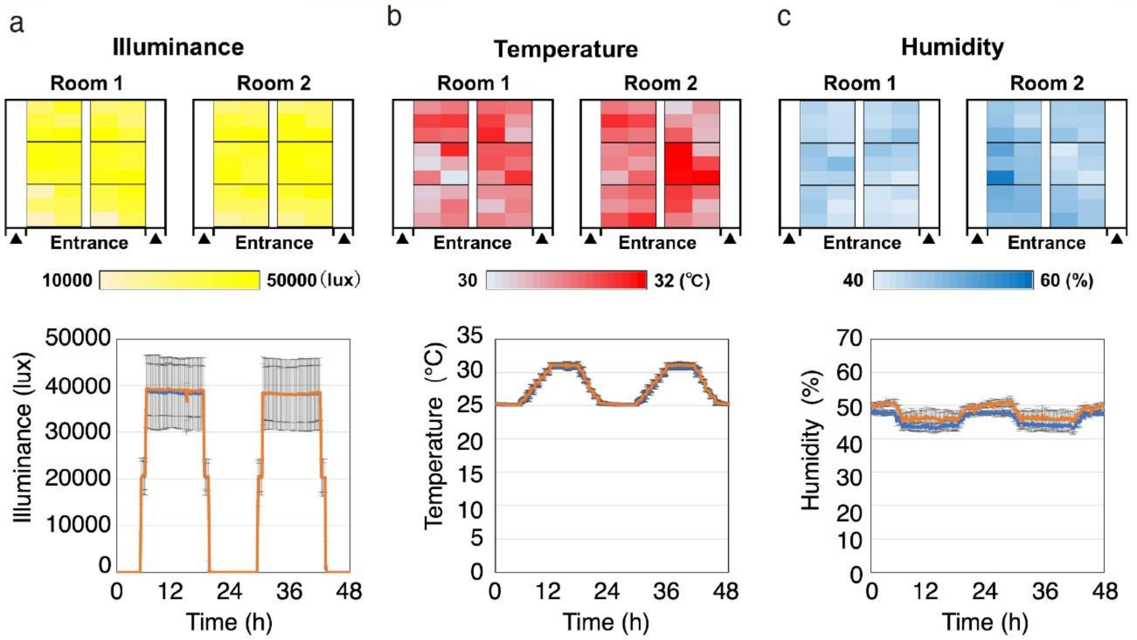


**Supplementary Figure 4. Environmental conditions in iPUPIL.**

**a–c.** Heatmap of light intensity (**a**), temperature (**b**), and humidity (**c**) observed across 72 pots in iPUPIL. Environmental variation within a 48-h period. The means ± SD of environmental parameters in six units in each room are shown.

### Supplementaly Figure 5

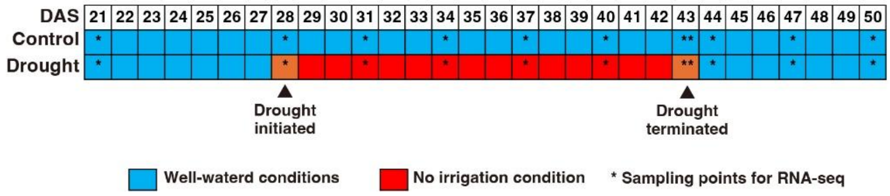


**Supplementary Figure 5. Experimental design for transcriptome analysis of rice plants in the control and drought plots.**

Rice plants were grown under control (well-watered) and drought (no irrigation) conditions. Sampling days for RNA-seq analyses are indicated. DAS: days after sowing.

#### Supplementary Figure 6

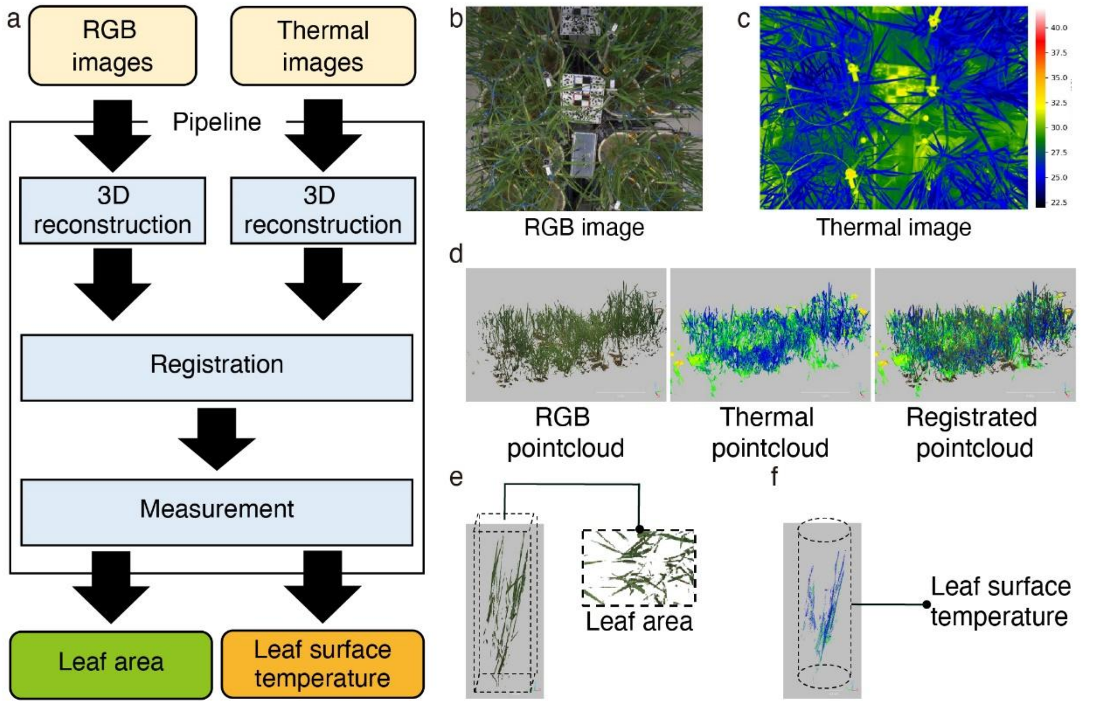


**Supplementary Figure 6. Overview of 3D image analysis of leaf area and temperature.**

**a.** The workflow of the 3D image analysis for RGB and thermal imagery. **b,c** Acquired original images. The resolution of the RGB image is 5,320 × 4,600 pixels (b). The resolution of the thermal image is 5,320 × 4,600 pixels (c). **d.** RGB, thermal, and registered point clouds are shown. **e.** The RGB point cloud was extracted as a cuboid region for each pot, and the area of the image projected from overhead is measured. **f.** The thermal point cloud was extracted as a cuboid region for each pot, and its average temperature was measured.

### Supplementaly Figure 7

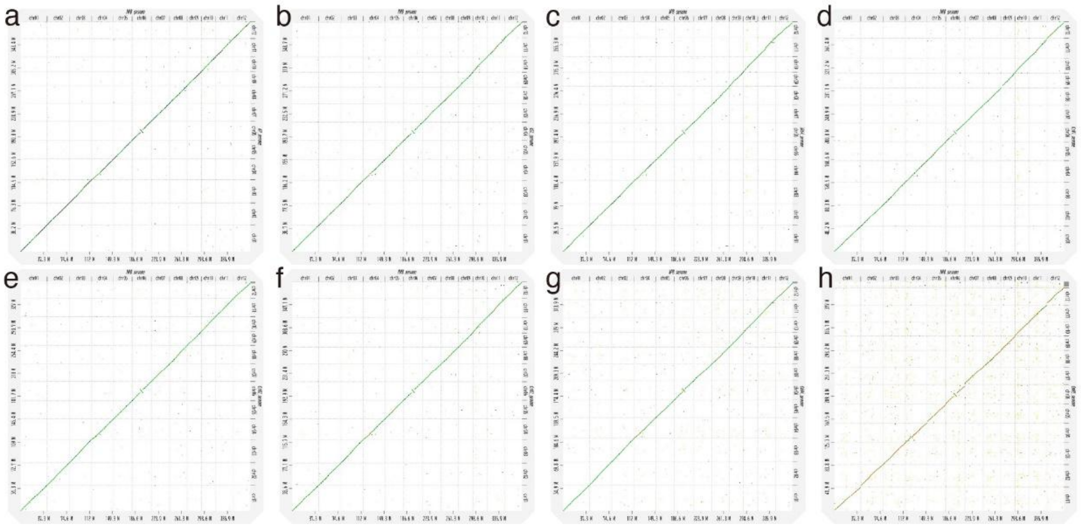


**Supplementary Figure 7. Dot plot of the Nipponbare genome and each assembled genome.**

**a**. Kinandang Patong (KP), **b**. Kasalath (KSL), **c**. IR64, **d**. OrW1, **e**. OrW2, **f**. OrW3, **g**. ObW4, **h**. OmW5. Two genomes were aligned using minimap2 (v2.28) with the option "-f 0.02" in the D-genies web application (v1.5.0). To reduce noise, the "Hide Noise" option was enabled.

#### Supplementaly Figure 8

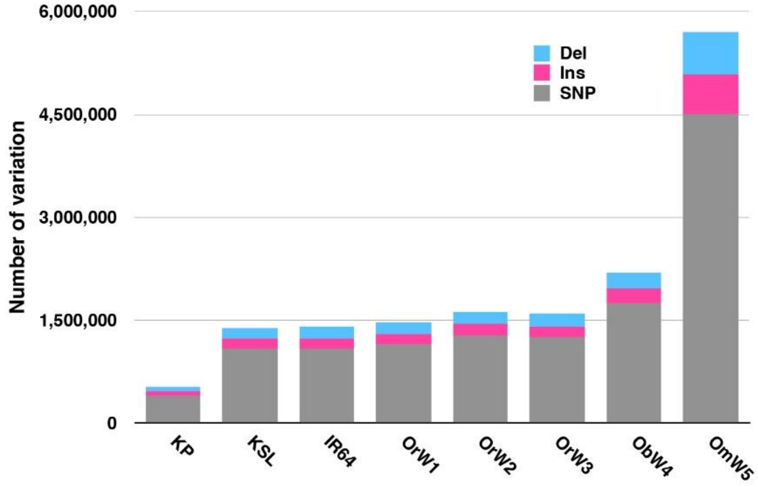


**Supplementary Figure 8. Number of single-nucleotide polymorphisms (SNPs) and structural variants (SVs) in the eight genomes compared with the Nipponbare reference genome.**

#### Supplementaly Figure 9

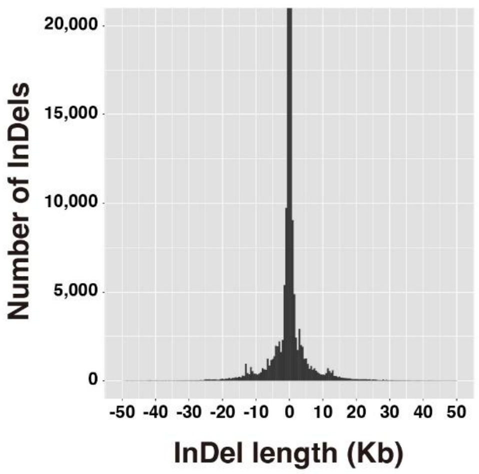


**Supplementary Figure 9. Length distribution of insertions and deletions (InDels) detected in the eight genomes compared with the Nipponbare reference genome.**

#### Supplementary Figure 10

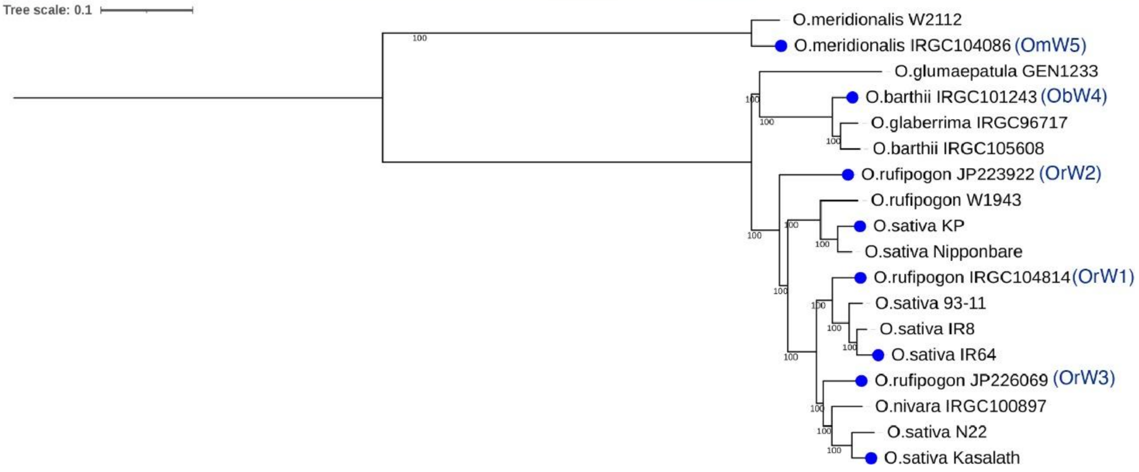


**Supplementary Figure 10. Phylogenetic tree of 18 rice genomes.**

A phylogenetic tree of the 18 rice accessions was constructed using the maximum likelihood method based on biallelic single-nucleotide polymorphism sites. The eight genomes established in this study were analyzed in conjunction with 10 previously reported genomes[1].

#### Supplementary Figure 11

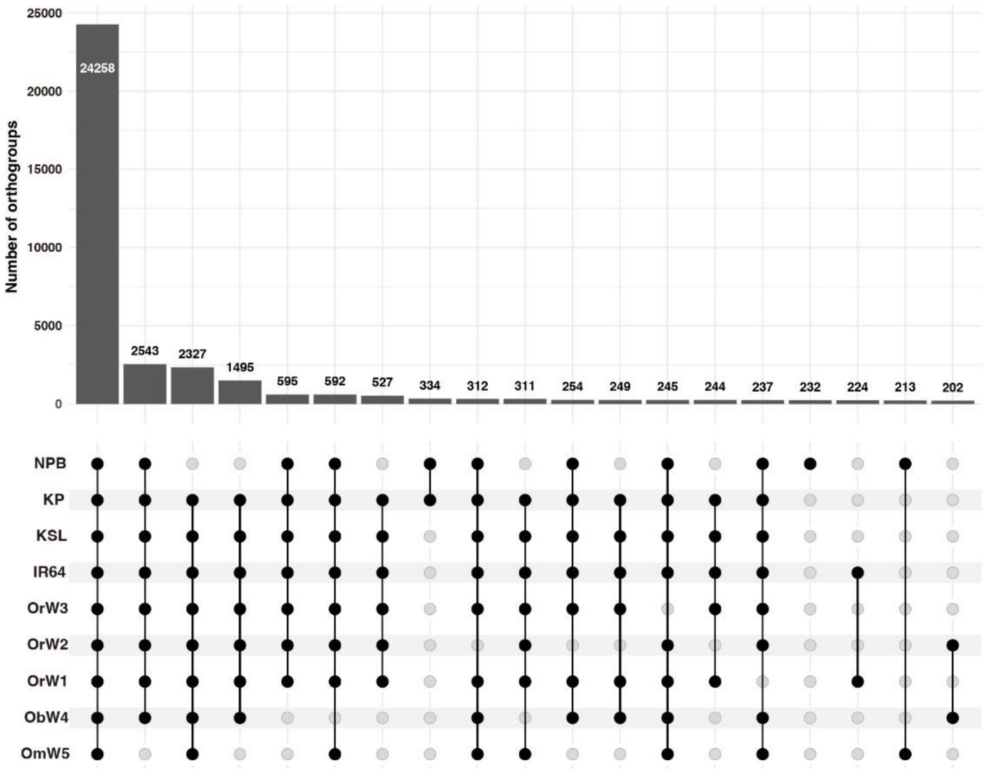


**Supplementary Figure 11. Orthogroup upset plot for the nine rice accessions.**

#### Supplementaly Figure 12

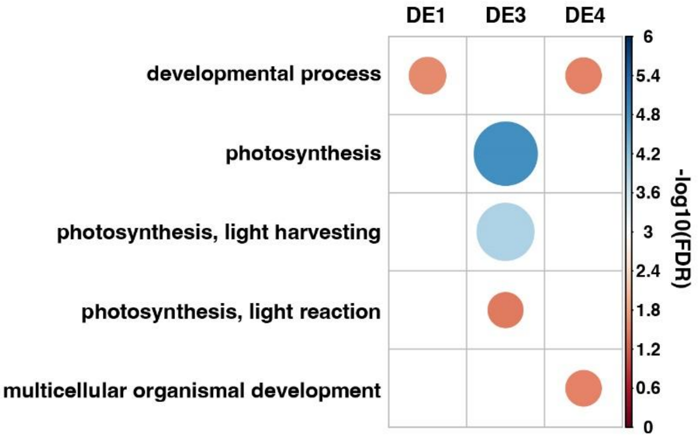


**Supplementary Figure 12. Enriched Gene Ontology (GO) terms in the "Biological process" category among differentially expressed genes in OrW2 compared with Nipponbare.**

Gene Ontology terms (Biological process) were enriched across each DEG cluster. FDR, false discovery rate. GO terms were not enriched in DE2.

#### Supplementary Figure 13

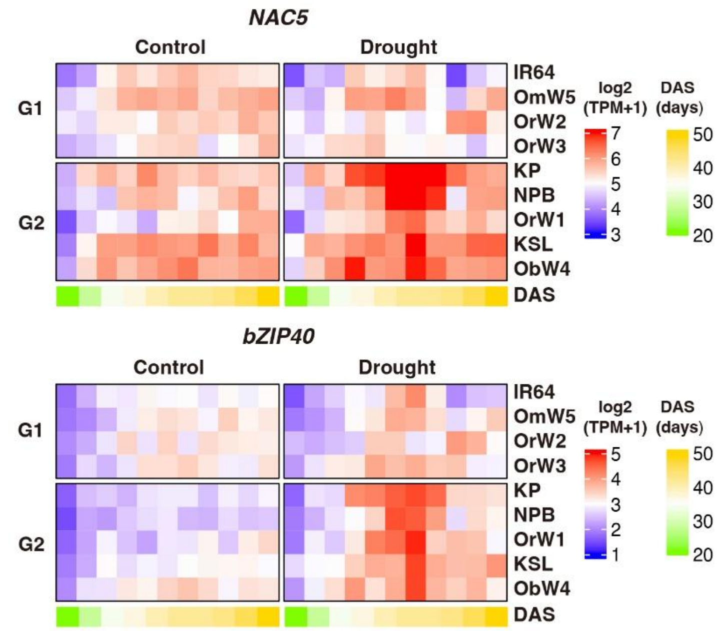


**Supplementary Figure 13. Expression patterns of genes encoding drought-inducible transcription factors.**

Heatmap of relative *NAC6* and *bZIP40* expression levels. DAS, days after sowing; NPB, Nipponbare; KP, Kinandang Patong; KSL, Kasalath.

#### Supplementary Figure 14

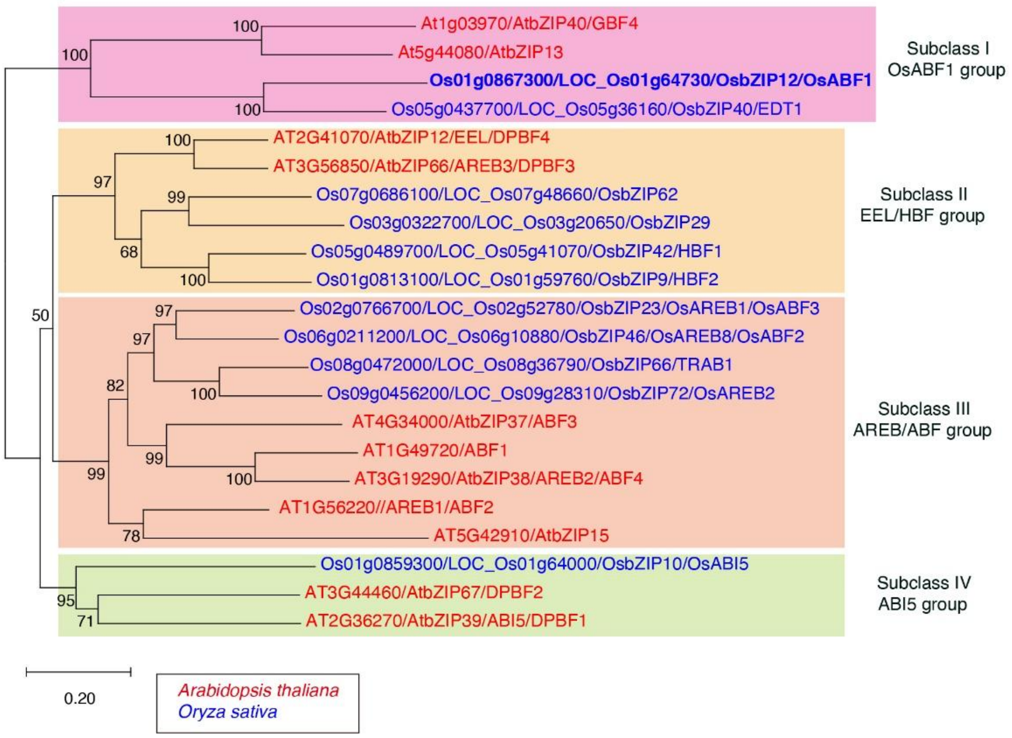


**Supplementary Figure 14. Phylogenetic analysis of ABF1 and its orthologous proteins.**

The phylogenetic tree was constructed from the full-length protein sequence of ABF1 and its orthologous proteins from *Arabidopsis thaliana* and *Oryza sativa* using the neighbor-joining method in MEGA11. The percentage of replicate trees clustered together in the bootstrap test (1,000 replicates) is shown next to the branches.

#### Supplementaly Figure 15

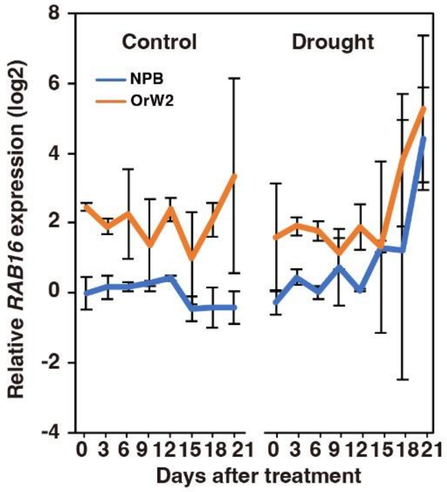


**Supplementary Figure 15. Expression patterns of the abscisic acid-mediated drought-responsive gene *RAB16* in Nipponbare and OrW2 in control and drought plots.**

The time course of relative *RAB16* expression levels is shown. Data represent the mean ± SD ($n$ = 3, biological replicates).

#### Supplementary Figure 16

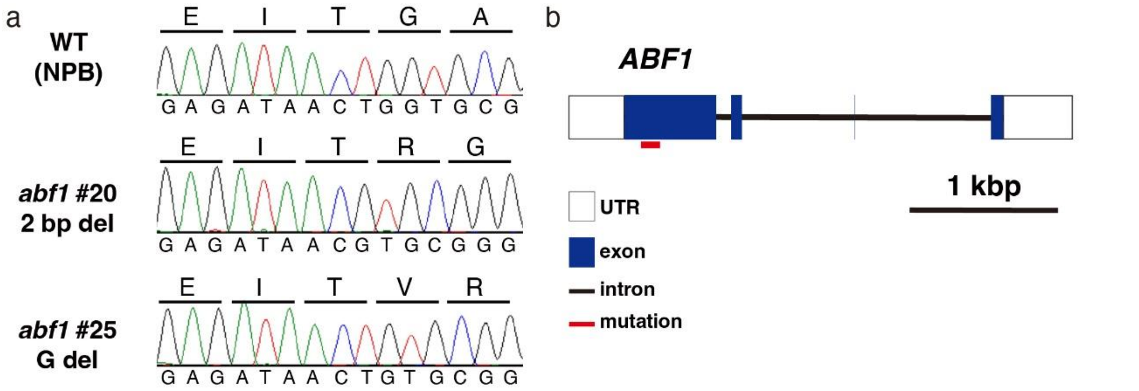


**Supplementary Figure 16. Loss-of-function *ABF1* mutants generated via CRISPR-Cas9 genome editing.**

**a.** CRISPR-Cas9-mediated mutation detected using Sanger sequencing. **b.** Mutations were detected in the first exon of *ABF1* in the mutants.

### Supplementary Figure 17

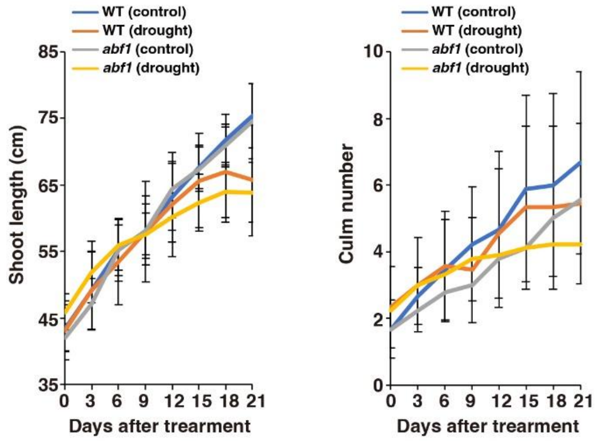


**Supplementary Figure 17. Growth of *abf1* mutant under drought stress.**

Water supply was stopped at 28 DAS (0 DAT) to 49 DAS (21 DAT) for the drought treatment. DAS, days after sowing; DAT, days after drought treatment. The time courses of shoot length and culm number per plant in the control and drought stress plots are shown. Data represent the mean ± SD ($n$ = 9).

#### Supplementary Figure 18

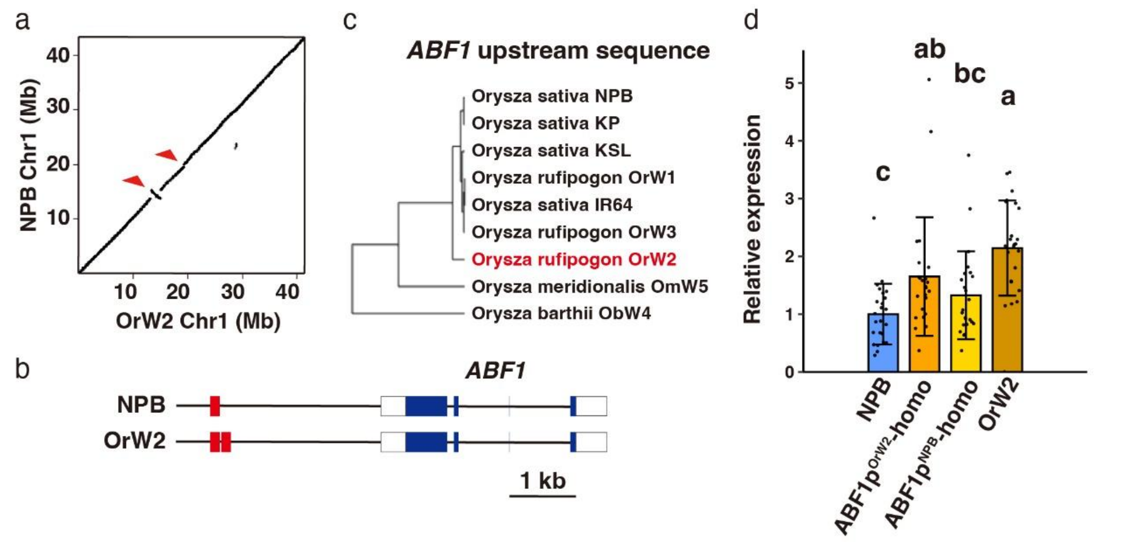


**Supplementary Figure 18. Single-nucleotide polymorphisms and structural variants around *ABF1* in OrW2.**

**a.** Genome alignment of chromosome 1 between Nipponbare (NPB) and Or2. Arrows indicate the alignment gap or inversion. The physical map position of *ABF1* is from 37,555,857 to 37,560,953 bp. **b.** Alignment of the upstream 3 kb sequence of *ABF1* transcription start sites. **c.** Schematic diagram of 3 kb upstream of the *ABF1* gene in NPB and OrW2. Part of the DNA sequence (red box) was duplicated in OrW2 compared with that in NPB. **d.** Relative *ABF1* expression levels in NPB and B2F2 lines under well-watered conditions.

#### Supplementary Figure 19

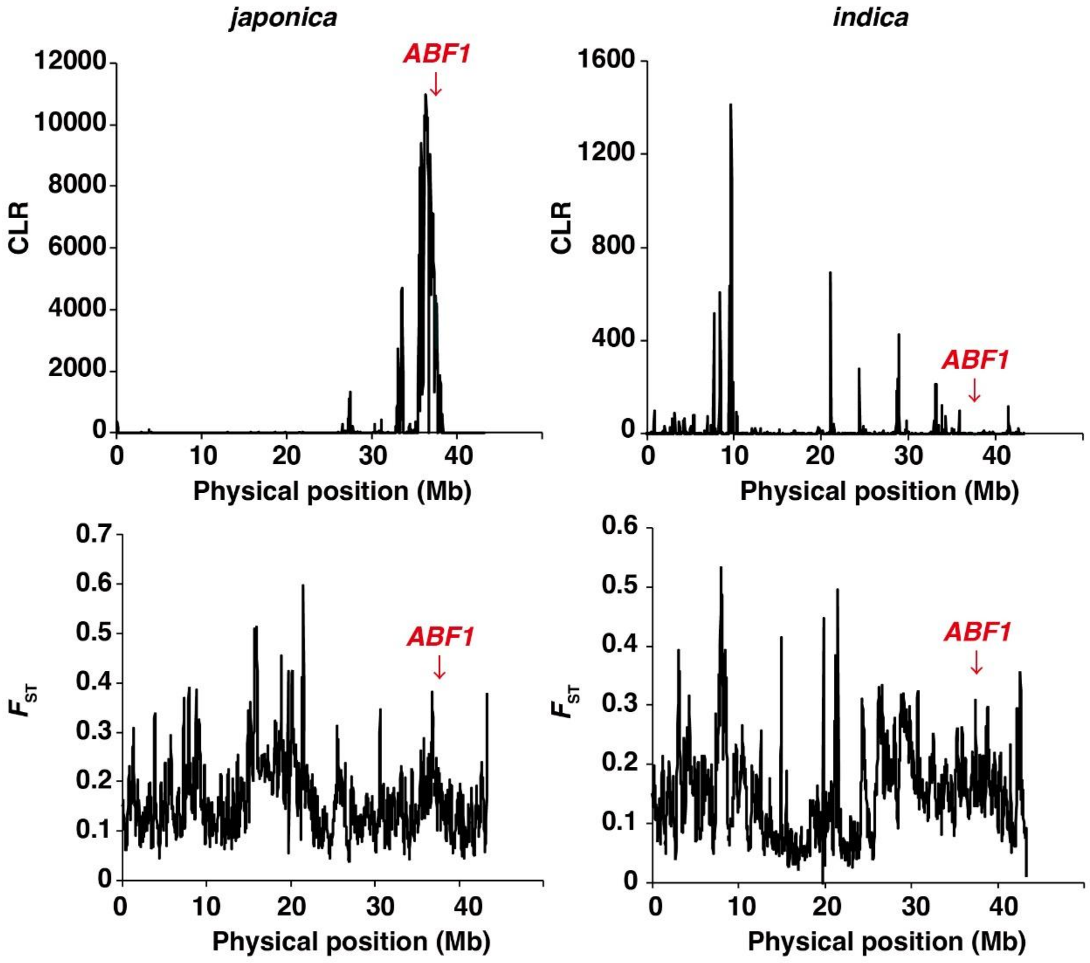


**Supplementary Figure 19. Detection of selective sweeps on rice chromosome 1.**

Physical position (Mb) and composite likelihood ratio (CLR) or fixation index ($F_{ST}$) are shown.

#### Supplementary Figure 20

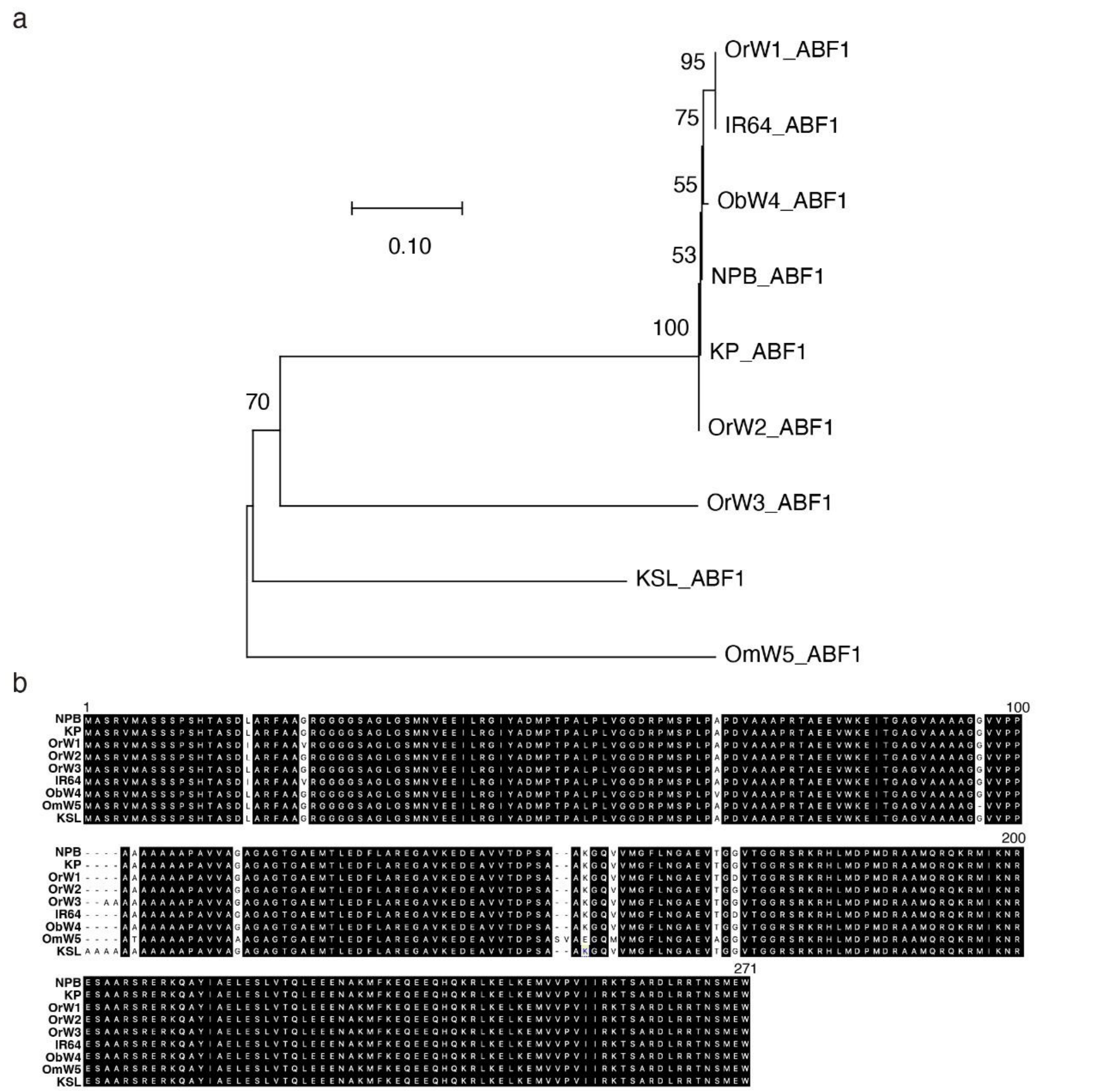


**Supplementary Figure 20. Variation in ABF1 protein sequences.**

**a.** Phylogenetic tree constructed from the full-length protein sequence of ABF1 using the neighbor-joining method in MEGA11. The percentage of replicate trees clustered together in the bootstrap test (1,000 replicates) is shown next to the branches. **b**. Alignment of ABF1 protein sequences.

#### Supplementary Figure 21

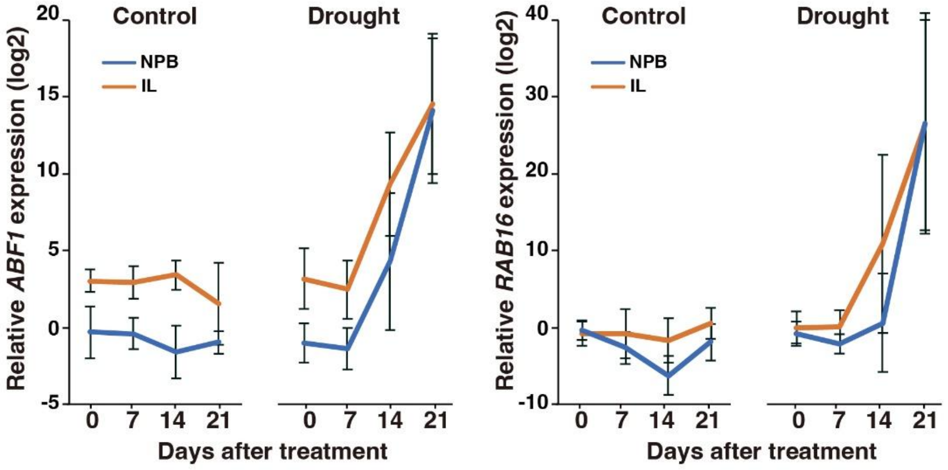


**Supplementary Figure 21. *ABF1* and *RAB16* expression patterns in introgression lines (ILs) under drought conditions.**

Water supply was stopped from 28 DAS (0 DAT) to 49 DAS (21 DAT) to induce drought stress. DAS, days after sowing; DAT, days after drought treatment. The time course of *ABF1* and *RAB16* expression in control and drought stress plots is shown. Data represent the mean ± SD ($n$ = 6).

#### Supplementary Figure 22

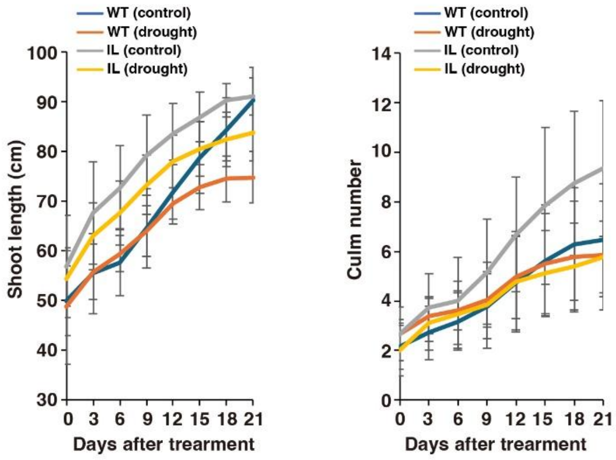


**Supplementary Figure 22. Growth of introgression lines (ILs) under drought conditions.**

Water supply was stopped from 28 DAS (0 DAT) to 49 DAS (21 DAT) to induce drought stress. DAS, days after sowing; DAT, days after drought treatment. Time courses of shoot length and culm number per plant in control and drought plots are shown. Data represent the mean ± SD ($n$ = 9).

#### Supplementary Figure 23

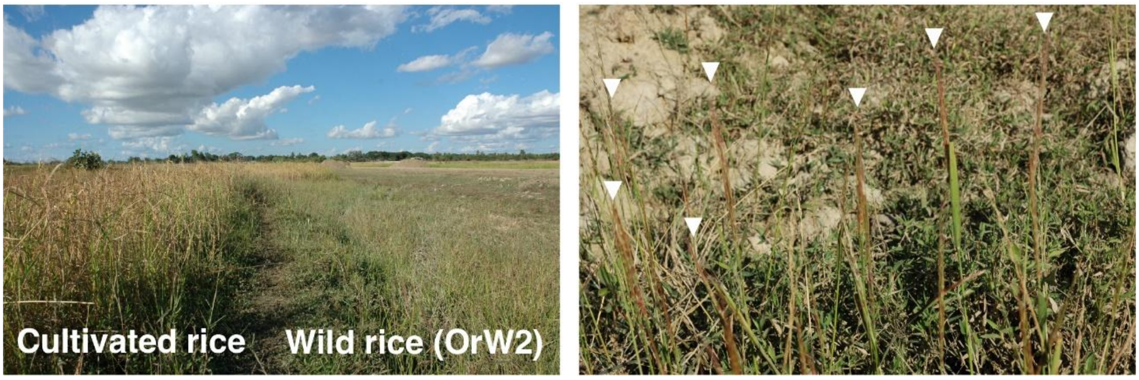


**Supplementary Figure 23. Native habitat of OrW2 in Myanmar.**

OrW2 seed collection near paddy fields in Rakhine State on November 29, 2004. Arrowheads indicate the panicles of OrW2.
